# Dual Histone Methyl Reader ZCWPW1 Facilitates Repair of Meiotic Double Strand Breaks

**DOI:** 10.1101/821603

**Authors:** Mohamed Mahgoub, Jacob Paiano, Melania Bruno, Wei Wu, Sarath Pathuri, Xing Zhang, Sherry Ralls, Xiaodong Cheng, Andre Nussenzweig, Todd Macfarlan

**Affiliations:** The Eunice Kennedy Shriver National Institutes of Child Health and Human Development, NIH, Bethesda, MD, USA; Laboratory of Genome Integrity, National Cancer Institute, NIH, Bethesda, MD, USA; Immunology Graduate Group, University of Pennsylvania, Philadelphia, PA, USA; Department of Epigenetics and Molecular Carcinogenesis, University of Texas MD Anderson Cancer Center, Houston, TX, USA

**Keywords:** ZCWPW1, PRDM9, Meiotic recombination, Double-Strand DNA Breaks, END-seq, CUT&RUN, Dual Histone Reader, Synapsis, Co-evolution

## Abstract

Meiotic crossovers result from homology-directed repair of double strand breaks (DSBs). Unlike yeast and plants, where DSBs are generated near gene promoters, in many vertebrates, DSBs are enriched at hotspots determined by the DNA binding activity of the rapidly evolving zinc finger array of PRDM9 (PR domain zinc finger protein 9). PRDM9 subsequently catalyzes tri-methylation of lysine 4 and lysine 36 of Histone H3 in nearby nucleosomes. Here, we identify the dual histone methylation reader ZCWPW1, which is tightly co-expressed during spermatogenesis with *Prdm9* and co-evolved with *Prdm9* in vertebrates, as an essential meiotic recombination factor required for efficient synapsis and repair of PRDM9-dependent DSBs. In sum, our results indicate that the evolution of a dual histone methylation writer/reader system in vertebrates facilitated a shift in genetic recombination away from a static pattern near genes towards a flexible pattern controlled by the rapidly evolving DNA binding activity of PRDM9.

## Introduction

Meiotic recombination (MR) generates genetic diversity in sexually reproducing organisms and facilitates proper synapsis and segregation of homologous chromosomes in gametes. During meiotic prophase I, recombination is initiated by programmed double strand breaks (DSBs) in DNA at thousands of specific 1-2 kb regions called hotspots (Kauppi et al., 2004). DSB initiation on one allele is necessary for homologous allele searching, pairing, and subsequent segregation. In many species, hotspots are distinguished by the presence of active histone marks in chromatin. For example, in yeast, plants and birds, hotspots are located at regions enriched with histone H3 tri-methylated on lysine 4 (H3K4me3), typically at gene promoters (Choi et al., 2013; Lam and Keeney, 2015; Lichten, 2015; Singhal et al., 2015). In contrast, in mammals, hotspots are determined by the DNA binding zinc finger array of PRDM9 (PR domain zinc finger protein 9)(Baudat et al., 2010; Myers et al., 2010; Parvanov et al., 2010). PRDM9 catalyzes tri-methylation of lysine 4 and lysine 36 of Histone 3 (H3K4me3 and H3K36me3 respectively) in nearby nucleosomes (Eram et al., 2014; Powers et al., 2016; Wu et al., 2013). This methyltransferase activity is essential for DSB formation (Diagouraga et al., 2018). *Prdm9* loss-of-function in *Mus musculus* leads to sterility (Hayashi et al., 2005), and many wild hybrid mice with incompatible *Prdm9* alleles are sterile (Davies et al., 2016), making *Prdm9* the only known speciation gene so far identified in mammals (Mihola et al., 2009). Furthermore, single nucleotide polymorphisms in PRDM9 have been linked to non-obstructive azoospermia in humans (Irie et al., 2009; Miyamoto et al., 2008)

During meiotic prophase I, chromatin re-organizes into condensed chromosomes, with DNA loops stemming from an axis composed of protein complexes (Cohen and Holloway, 2014; Xu et al., 2019). MR occurs simultaneously with homologous chromosome pairing and synapsis, and the relationship between these events is complex (Santos, 1999). Recombination is achieved via homologous repair of programmed DSBs at hotspots. To initiate DSBs, DNA loops at which hotspots are located must be tethered to the chromosomal axis for recruitment to SPO11, the type II topoisomerase responsible for generating DSBs (Keeney, 2008; Lange et al., 2016). After inducing a DSB, DNA ends are resected and single strand DNA binding proteins DMC1 and RAD51 are recruited to facilitate homologous strand invasion, formation of Holliday junctions, and subsequent resolution as either a crossover or non-crossover (Gray and Cohen, 2016). Synapsis is achieved simultaneously with MR by connecting the two proteinaceous cores of each homolog axis through a central region containing the synaptic protein SYCP1, which self assembles via its N-terminus, facilitating the closure of the synaptonemal complex (SC) like a zipper (Cohen and Holloway, 2014). In mice, a fully assembled SC is required for later steps in recombination (Hamer et al., 2008; Stack, 1984). Thus, the formation of the SC may facilitate MR by physically connecting the two chromosomes. Likewise, SC formation and proper synapsis requires MR machinery including Dmc1, Rad51, and Mre11 (Rockmill B et al, Genes Dev 1995, Nairz K. et al Genes Dev 1997).

Despite PRDM9’s role in specifying hotspots in mice, DSBs are still produced in *Prdm9* knock-out (KO) mice, but they are re-positioned to the “default” position at promoters (Brick et al., 2012). However, these DSBs are not repaired efficiently, leading to meiotic arrest and partial asynapsis. In F1 hybrid mice, the presence of PRDM9 binding motifs on both the cut and uncut homolog improves recombination rates (Hinch et al., 2019; Li et al., 2019b), suggesting that PRDM9 itself, and perhaps its downstream factors, play important roles beyond merely initiating DSBs.

*Prdm9* first evolved in jawless vertebrates, and it has been either completely or partially lost in several lineages of vertebrates, including several fish lineages, in birds and crocodiles, and in canids, indicating it is not absolutely required for MR, synapsis and fertility (Baker et al., 2017). Furthermore, *Prdm9* KO mice crossed into other strain backgrounds partially restores fertility in male mice (Mihola et al., 2019). Nonetheless, the evolution of PRDM9 in vertebrates replaced a pre-existing hotspot selection system based on the presence of single H3K4me3 marks at promoters with a new selection system based on the presence of dual H3K4me3/H3K36me3 marks at PRDM9 binding sites that facilitated more rapid DSB repair and synapsis. In yeast, the histone reader Spp1 links H3K4me3 sites at promoters with the MR machinery, promoting DSB formation (Adam et al., 2018). However in mice, the Spp1 orthologue CXXC1, which also interacts with PRDM9, does not play a role DSB generation or MR (Tian et al., 2018). We reasoned other meiotic factors might have evolved in vertebrates to link PRDM9 to either the MR or SC machinery, that would interact directly with the histone marks deposited by PRDM9. Here, we identify Zinc Finger CW-Type and PWWP Domain Containing 1 (ZCWPW1) as a dual histone methylation reader specific for PRDM9 catalyzed histone marks (H3K4me3 and H3K36me3) that facilitates the repair of PRDM9-induced DSBs. Our study reveals a novel histone methylation reader/writer system that controls patterns of meiotic recombination in mammals.

## Results

### ZCWPW1 is a dual histone methylation reader co-expressed with PRDM9 in spermatocytes

To identify PRDM9 co-factors that may play a role in meiotic recombination, we searched for *Prdm9* co-expressed genes in single cells during meiosis from a published dataset (Chen et al., 2018). As expected, the top *Prdm9*-correlated genes are known factors in spermatogenesis or DNA metabolism (Figure 1A, Table S1), however the most correlated gene in our analysis was *Zcwpw1* (rho = 0.519, p-value = 2E-6). Similar to *Prdm9*, *Zcwpw1* mRNA is highly expressed exclusively in the testis in both mouse and human (Figure S1) and ZCWPW1 protein expression is mostly restricted to the testis in mice (Figure 1B). ZCWPW1 has three recognizable domains: SCP1, zf-CW and PWWP (Figure 1C). The PWWP domain found in multiple proteins binds specifically to histone H3 containing the H3K36me3 mark (Qin and Min, 2014), while the zf-CW domain of ZCWPW1 was found to possess H3K4me3-specific binding (He et al., 2010). The SCP1 domain has homology to the synaptonemal complex protein 1 (SYCP1), the major component of the transverse filaments of the synaptonemal complex (De Vries et al., 2005). The N-terminal region of SYCP1 homologous to the SCP1 domain of ZCWPW1 plays a role in SYCP1 dimerization that facilitates synapse assembly (Seo et al., 2016).

**Figure 1:**
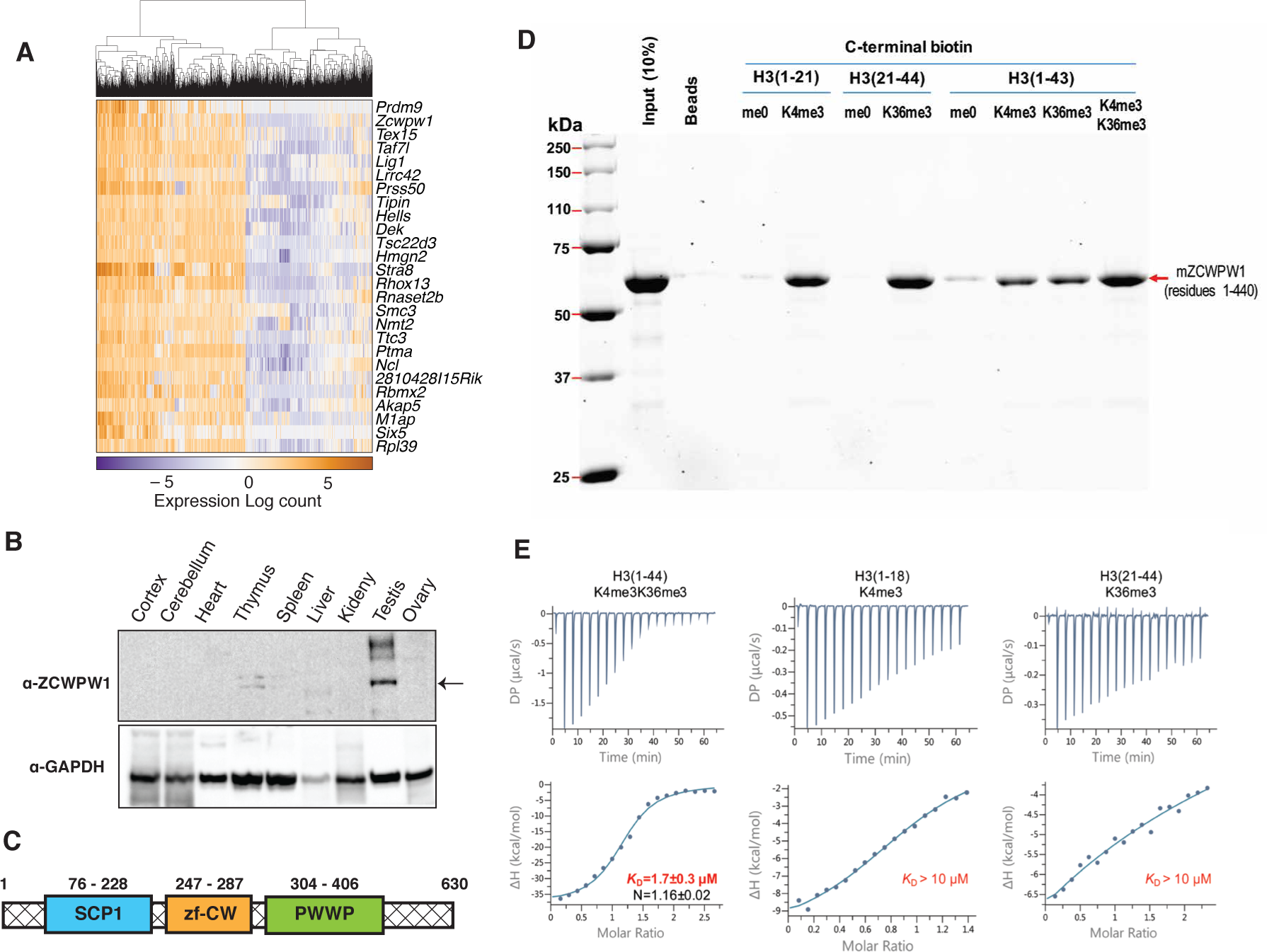
ZCWPW1 tissue expression and methyl histone binding activity *in vitro*. (A) Heatmap of the top 25 genes co-expressed with PRDM9 during meiosis ordered by PRDM9 correlation coefficient (*rho*). (B) Western blot with indicated antibodies in different tissues from WT B6 mice. The arrow indicates ZCWPW1 band. (C) Mouse ZCWPW1 protein with structural domains shown as annotated from NCBI domain database. (D) Recombinant ZCWPW1 was mixed with indicated biotinylated histone peptides and bound proteins were identified by Coomassie staining. (E) Isothermal calorimetry measurements were performed with recombinant ZCWPW1 and indicated histone peptides. *KD* and stoichiometry values are displayed.

Since PRDM9 catalyzes the formation of both H3K4me3 (Hayashi et al., 2005; Wu et al., 2013) and H3K36me3 (Eram et al., 2014; Powers et al., 2016), we reasoned that ZCWPW1 may link the PRDM9-induced histone marks to the meiotic recombination machinery by binding to the dually marked histone tails. Using *in vitro* biotin-streptavidin pulldown assays, we determined that recombinant ZCWPW1 (residues 1-440) binds with higher affinity to H3K4me3 and H3K36me3 than non-methylated biotinylated H3 peptides (Figure 1D). Importantly, dual-modified H3K4me3/K36me3 peptides had the highest binding affinity for ZCWPW1 compared to peptides with either single modification (Figure 1D). We further quantified the binding of ZCWPW1 to histone peptides by Isothermal Titration Calorimetry, which demonstrated that ZCWPW1 binds with the highest affinity to H3K4me3/K36me3 peptides (*KD* =1.7±0.3 µM) compared to H3K4me3 and H3K36me3 alone (*KD* > 10 µM each), with a 1:1 stoichiometry (Figure 1E). In sum these results indicate that ZCWPW1 is a meiotic histone methylation specific reader protein for the unique dual methyl marks catalyzed by PRDM9, suggesting that ZCWPW1 and PRDM9 have complementary roles in meiosis.

### Co-evolution of *Zcwpw1* and *Prdm9* in Vertebrates

PRDM9 has an extraordinary evolutionary pattern as it possesses a rapidly evolving zinc finger array leading to distinct hotspots even among species belonging to the same genus (Baker et al., 2017; Oliver et al., 2009; Thomas et al., 2009). Furthermore, while it emerged in jawless fishes, it has repeatedly been lost in several species across different clades including in birds, crocodiles, and amphibians, as well as in several lineages of fish. To determine whether *Prdm9* and *Zcwpw1* share a similar evolutionary pattern, we retrieved all *Zcwpw1* orthologs from both OrthoDB (Kriventseva et al., 2019) and Ensembl databases (Table S2). Multiple sequence alignment demonstrated conservation of SCP1, zf-CW and PWWP domains (Figures 2A and S2). Unlike zf-CW and PWWP domains, which are highly conserved among all orthologs, the SCP1 domain showed higher conservation amongst turtle and mammal orthologs (Figure S2; Document S1). To compare the domain architecture of ZCWPW1 orthologs with the previously described PRDM9 orthologs across different clades (Baker et al., 2017), we used the PrositeScan tool (de Castro et al., 2006) to screen for zf-CW and PWWP domains in all orthologs of ZCWPW1 (Table S2). The SCP1 domain was excluded because it doesn’t have any matching domain in that database. Although the zf-CW and PWWP domains can be found throughout metazoans, ZCWPW1-like proteins containing both domains in tandem first appear in jawless vertebrates similar to PRDM9 (Figure 2B). Furthermore, ZCWPW1 orthologs have been completely lost in birds and crocodiles. PRDM9 orthologs in at least one monotreme (Platypus) and one marsupial (Tasmanian devil) have a truncation in the KRAB domain, which is speculated to prevent PRDM9 from playing a role in meiotic recombination (Imai et al., 2017), and some species in these clades (Platypus and Opossum), although containing a ZCWPW1-like ortholog, lack the PWWP domain. This striking co-evolutionary pattern of PRDM9 and ZCWPW1 implies that full or partial loss of either factor may facilitate the loss of the other, and that the two genes are essential in a cooperative manner.

**Figure 2:**
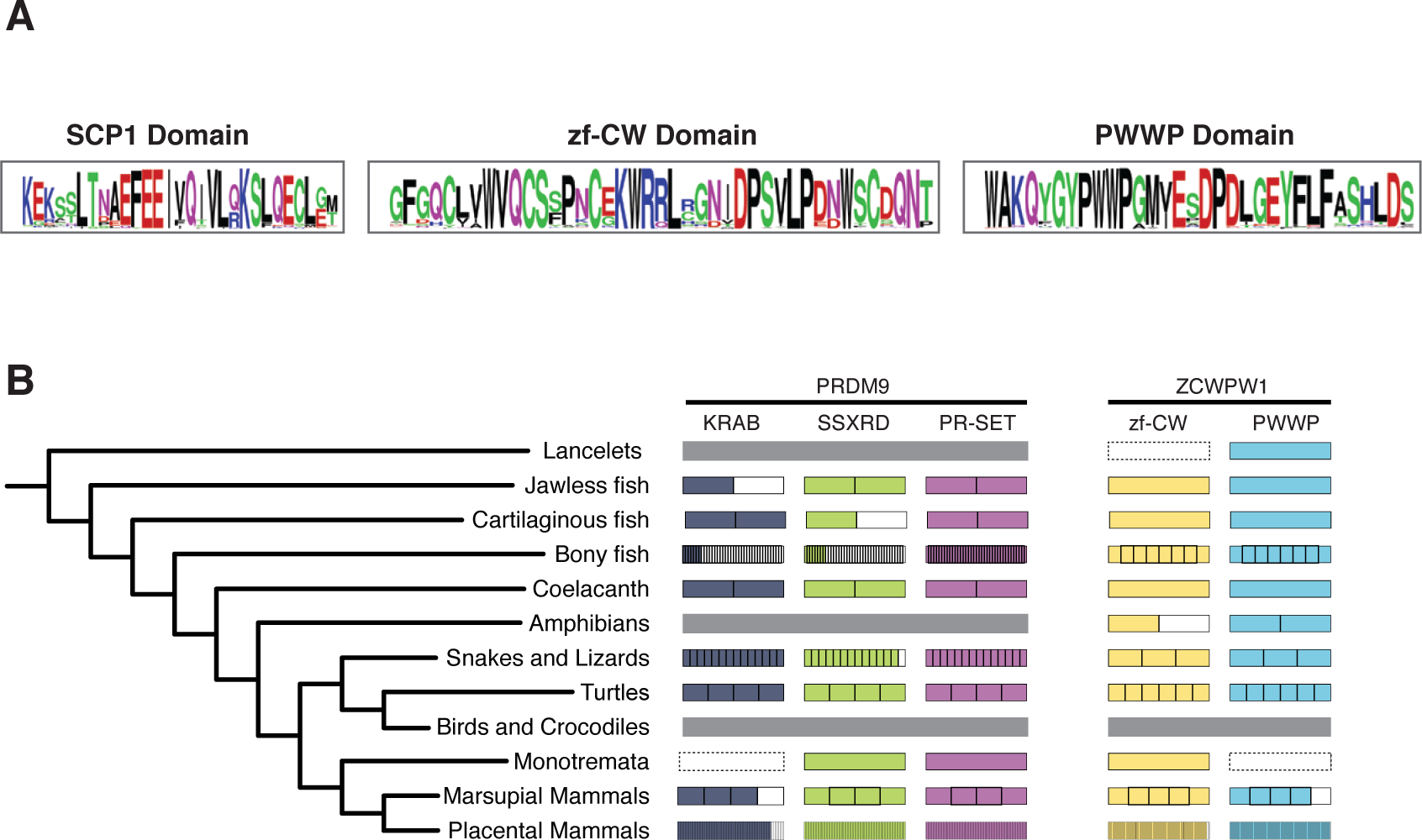
Co-evolution of *Zcwpw1* and *Prdm9* in vertebrates. (A) Sequence logos for ZCWPW1 consensus motifs. ZCWPW1 orthologs were aligned and the region corresponding to each motif was extracted for logo generation. (B) Domain structure and conservation for ZCWPW1 and PRDM9 orthologs across clades. The N-terminal portion of PRDM9 domain structure is reproduced from (Baker et al. 2017). Amino acid sequences of ZCWPW1 orthologs were retrieved and combined from OrthoDB and Ensembl databases, and each ortholog’s domain structure was determined by scanning for the presence of zf-CW and PWWP domains using ScanProsite tool, then all orthologs were grouped according to clades. For each clade, a protein domain is represented by a large rectangle subdivided into smaller sub-rectangles, in accordance with the number of screened species per clade, with color indicating conservation of the domain in the single species. Dashed border means no species in the clade had the domain of interest, and large grey rectangle indicates lack of the whole protein in all species examined in that clade.

### ZCWPW1 binds specifically to dual methylated nucleosomes at meiotic hotspots

Meiotic recombination hotspots in mice are primarily determined by PRDM9 binding (Baudat et al., 2010; Davies et al., 2016; Grey et al., 2017; Myers et al., 2010; Parvanov et al., 2010). These hotspots are characterized by PRDM9-dependent histone H3 methylation (H3K4me3 and H3K36me3) (Baker et al., 2014; Grey et al., 2011, 2018; Hayashi et al., 2005; Smagulova et al., 2011). To determine whether ZCWPW1 binds to these chromatin sites *in vivo*, we performed Cleavage Under Targets and Release Using Nuclease (CUT&RUN) (Christoph and Siegenthaler, 2016) using custom polyclonal ZCWPW1 antibodies on mouse spermatocytes. We identified 4,300 ZCWPW1 peaks (*p* < 0.001). Most ZCWPW1 peaks overlap with previously identified hotspots defined by the presence of the single-strand (ss) DNA binding protein DMC1 (Brick et al., 2018) (Figure 3A) or released SPO11 oligos (Lange et al., 2016) (Figure S3A) (82.4% and 84.2% respectively). PRDM9 binding generates a nucleosome-depleted region with nucleosomes organized symmetrically around PRDM9 binding sites (Baker et al., 2014). ZCWPW1 and H3K4me3 mapping using CUT&RUN showed a similar nucleosome distribution at hotspots determined by DMC1 binding (Figure 3A) and SPO11 oligos (Figure S3A). It is notable that DSB hotspots that fail to overlap with ZCWPW1 peaks still show ZCWPW1 signal that is below the peak detection threshold (Figure S3B). Interestingly, ZCWPW1 does not occupy gene promoters which contain abundant H3K4me3 marks, with only 0.9% of ZCWPW1 peaks overlapping transcription starting sites (Figure 3B). This suggests - in agreement with *in vitro* binding assays (Figures 1D and 1E) – that the presence of H3K4me3 modification alone is not sufficient for efficient recruitment of ZCWPW1 to nucleosomes in spermatocytes. To investigate whether the presence of both H3K4me3 and H3K36me3 marks are necessary for ZCWPW1 recruitment to chromatin *in vivo*, we used a published ChIP-seq dataset of H3K4me3 and H3K36me3 binding in prophase I spermatocytes (Lam et al., 2019). The vast majority of ZCWPW1 peaks overlapped with regions containing both H3K4me3 and H3K36me3 marks (85%), with significantly less overlapping either mark alone (Figure 3C and 3D). This demonstrates that ZCWPW1 binding is determined by the concurrent presence of dual H3K4me3/H3K36me3 marks, the same modifications generated by PRDM9.

**Figure 3:**
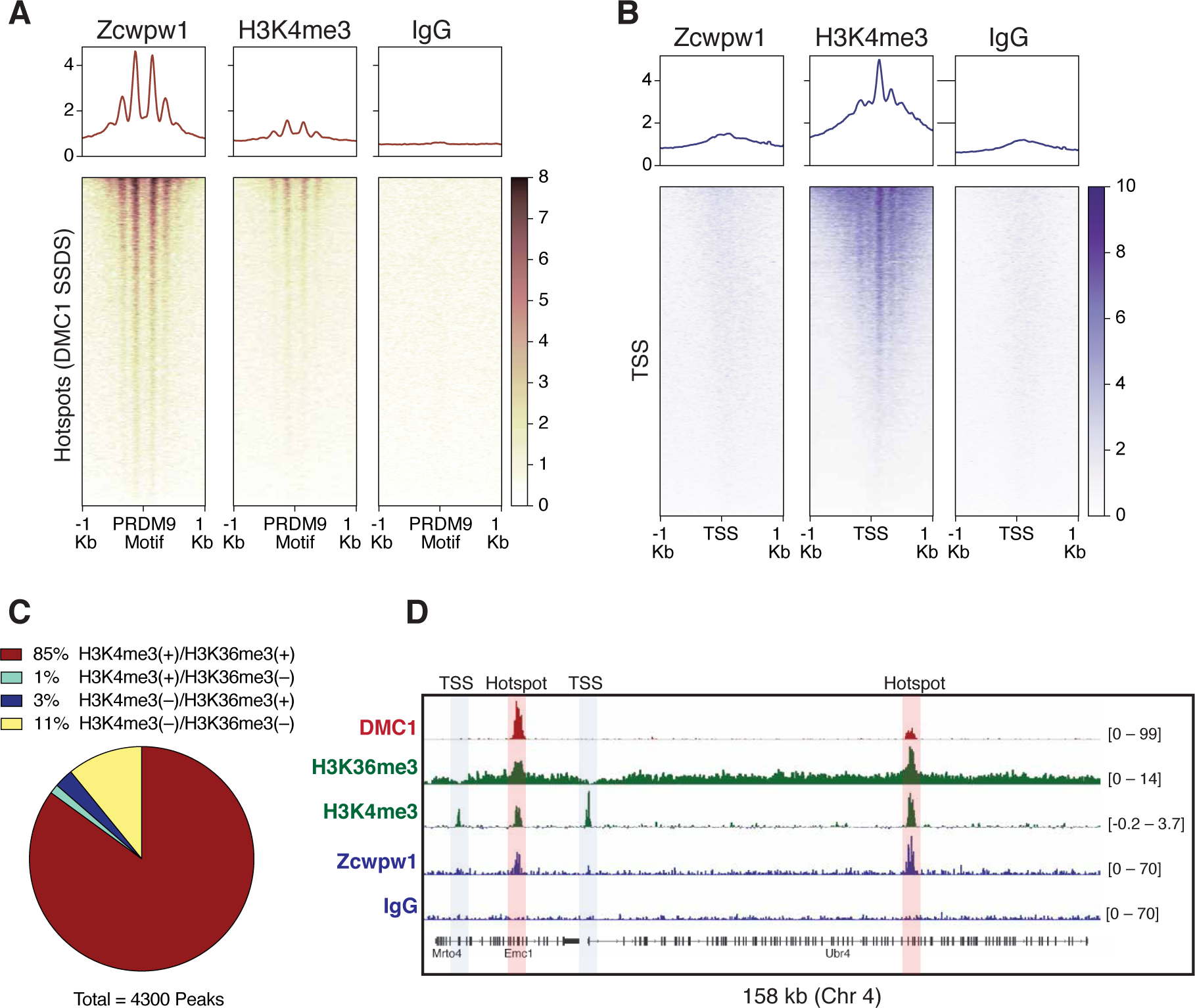
Mapping of ZCWPW1 chromatin biding *in vivo* using CUT&RUN in spermatocytes from B6/B6 mice. (A) Heatmaps representing ZCWPW1, H3K4me3 and IgG (anti-GFP) occupancy in B6/B6 mouse spermatocytes at hotspots determined by DMC1 SSDS (GSE99921). Signals are centered around PRDM9 motifs. Hotspots with multiple motifs were excluded from plotting. (B) ZCWPW1, H3K4me3 and IgG (anti-GFP) occupancy in B6/B6 spermatocytes at transcription starting site (TSS). (C) Pie chart showing methylation states of histone H3 at ZCWPW1 peaks in B6/B6 spermatocytes. (D) Read coverage plots for DMC1 SSDS hotspots (GSE35498/red), H3K4me3/H3K36me3 (GSE121760/Green) and ZCWPW1/IgG (CUT&RUN/blue) across a region on chromosome 4. Hotspots and TSSs are highlighted.

### ZCWPW1 chromatin occupancy is determined by PRDM9 in allele specific manner

To determine factors responsible for ZCWPW1 recruitment to chromatin *in vivo* in an unbiased manner, we searched for enriched motifs within ZCWPW1 peak regions. This analysis identified exclusively the PRDM9 binding motif of the Dom2 allele (PRDM9^Dom2^), found within the C57Bl/6J strain (*p* = 1e-261) (Figure S3C; Table S3), suggesting that ZCWPW1 occupancy is solely PRDM9-dependent. To test this hypothesis, we mapped ZCWPW1 binding in F1-hybrid mouse spermatocytes from a C57Bl/6J (B6) and CAST/EiJ (CAST) cross, since CAST mice encode a unique PRDM9 allele (PRDM9^Cast^) with distinct DNA binding properties. In these mice, we identified 11,027 ZCWPW1 peaks, more than twice the number of peaks in pure B6/B6 mice. 62% of these peaks overlap with previously described CAST/CAST hotspots (Smagulova et al., 2016), while only 18.1% overlap with B6/B6 hotspots (Figure S4A), a finding consistent with reported PRDM9^Cast^ dominance in hybrid mice (Baker et al., 2015). Furthermore, ZCWPW1 also binds to 1,892 novel hotspots that do not exist in either B6/B6 or CAST/CAST mice, but only appear in F1-hybrids, where PRDM9^Cast^ binds to the B6 genome and PRDM9^Dom2^ binds to the CAST genome. These sites have been shown to be due to PRDM9-binding-site loss that occurs as a result of accumulated gene conversion events within each sub-species (Baker et al., 2015). De novo motif discovery in the hybrids revealed two motifs identical to PRDM9^Cast^ and PRDM9^Dom2^ (*p* = 1e-768 and 1e-60 respectively) (Figure 4A; Document S2). This resemblance between ZCWPW1 and PRDM9 in their chromatin binding motifs is reflected in the pattern of nucleosome occupancy of ZCWPW1 genome wide in a PRDM9-allele specific manner in both B6/B6 and B6/CAST mice (Figures 4B and S4B). Overall, these results indicate that ZCWPW1 occupancy is a novel marker for meiotic hotspots that reflects PRDM9 allele specificity.

**Figure 4:**
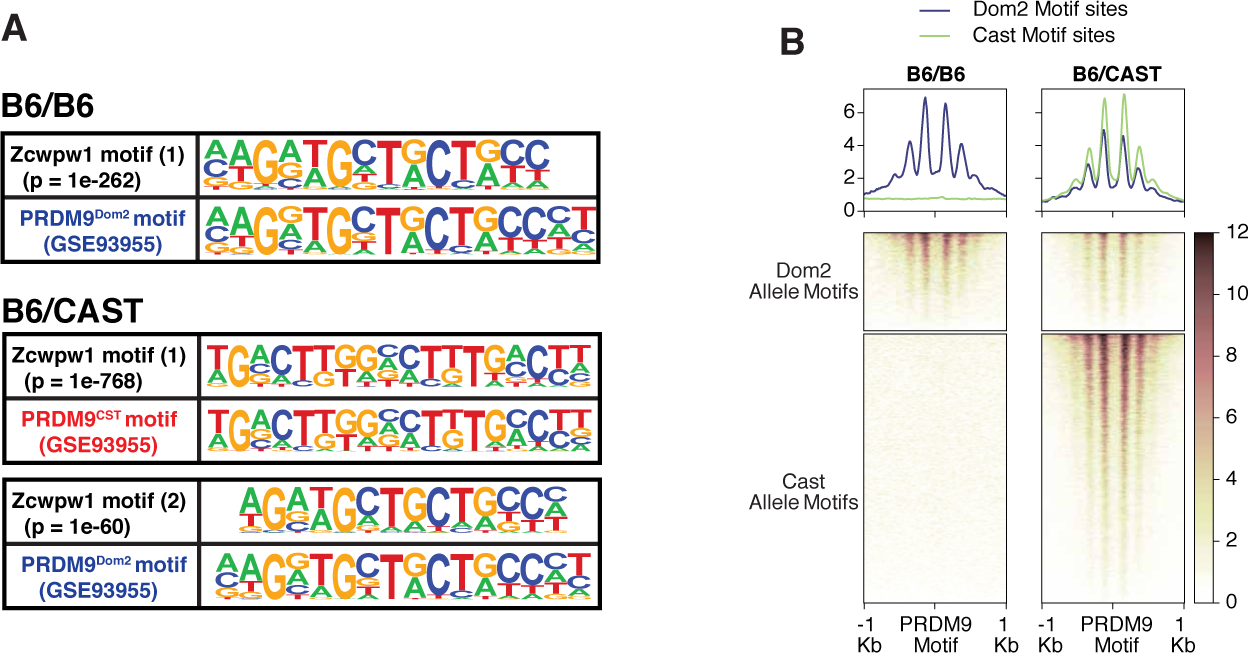
Mapping of ZCWPW1 chromatin biding *in vivo* using CUT&RUN in spermatocytes from F1 B6/CAST hybrid mice. (A) Comparisons of de novo discovered motifs for PRDM9^Dom2^, PRDM9^Cast^ and ZCWPW1. For PRDM9^Dom2^ and PRDM9^Cast^, ChIP-seq peaks (GSE93955) for either allele were queried by HOMER to identify the consensus DNA motif for their binding sites. For ZCWPW1, CUT&RUN peaks from either B6/B6 or B6/CAST F1 hybrid were used to identify ZCWPW1 binding motifs in pure and mixed genomic backgrounds, respectively. The first motif hit is shown for binding in B6/B6 and the first and second hits are shown for B6/CAST. (B) Heatmaps comparing ZCWPW1 occupancy in spermatocytes from either B6/B6 or B6/CAST F1 hybrid. ZCWPW1 signal is plotted around F1 hybrid hotspots (GSE73833) in mm10 genome assembly. Regions are categorized according to PRDM9 allele motifs (*Prdm9^Dom2^* and *Prdm9^Cast^* for B6 and CAST strains respectively). Signals are centered around PRDM9 motifs.

### *Zcwpw*1 knockout mice are azoospermic and display asynapsis and repair defects

PRDM9 induced DSBs are critical for successful chromosome synapsis during meiosis, and *Prdm9*-null male mice (*M. m. domesticus)* are azoospermic as a result of meiotic arrest due to compromised DSB repair and chromosomal asynapsis (Hayashi et al., 2005). As ZCWPW1 binding overlapped PRDM9 binding sites, we hypothesized it also plays important role in homolog synapsis during meiosis. To test that hypothesis, we generated mice with homozygous deletion of *Zcwpw1* (*Zcwpw1^KO/KO^*), which were born healthy and at mendelian ratios. Western blotting confirmed the absence of ZCWPW1 in testis from *Zcwpw1^KO/KO^* mice (Figure S5A), which have significantly smaller testes (80 mg) compared to wild type (*Zcwpw1^WT/WT^*) (191 mg) (Figure S5B). Hematoxylin Eosin (H&E) staining of tissue sections demonstrated the total absence of spermatids in the seminiferous tubules of *Zcwpw1^KO/KO^* mice testes (Figures 5A and S5C). Double immunofluorescence staining of chromosome spreads from spermatocytes with the SC axial element marker SYCP3 and the SC central element marker SYCP1 revealed *Zcwpw1^KO/KO^* mice display partial chromosomal asynapsis and arrest at pachytene-like stage of prophase I (Figure 5B), consistent with a recent report (Li et al., 2019a).

**Figure 5:**
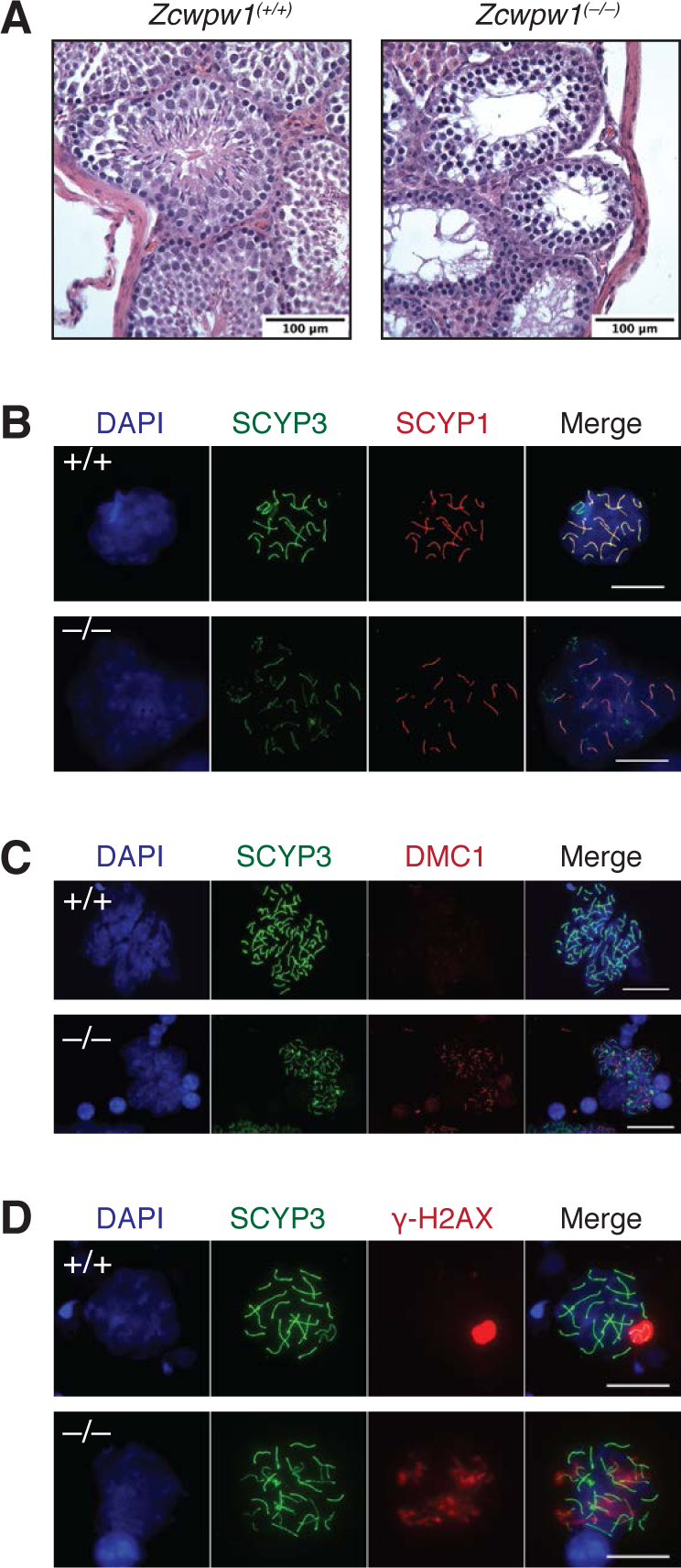
*Zcwpw1^KO/KO^* mice are azoospermic and display hallmarks of defective synapsis and DSB repair. (A) Hemataoxylin & Eosin staining of paraffin embedded tissue sections from testes of *Zcwpw1^WT/WT^* or *Zcwpw1 ^KO/KO^*. Scale bar is 100 µm. (B-D) Indirect immunofluorescence of spermatocyte spreads from *Zcwpw1^WT/WT^* or *Zcwpw1^KO/KO^* with indicated antibodies. Scale bar is 10 µm.

As synapsis is initiated by PRDM9-induced DSBs, we tested whether asynapsis in spermatocytes from *Zcwpw1^KO/KO^* mice were due to failure in DSB formation. We stained spermatocytes with antibodies recognizing the DSB marker γ-H2AX and the repair factor DMC1, which is recruited to single stranded DNA resulting from DSBs. Spermatocytes from *Zcwpw1^KO/KO^* mice have widespread focal accumulation of γ-H2AX and DMC1 in arrested cells, unlike pachytene stage cells in *Zcwpw1^WT/WT^* mice, in which efficient repair prevents such accumulation (Figures 5C and 5D). These results suggest intact DSB formation by SPO11 but defective repair and synapsis. Collectively, *Zcwpw1^KO/KO^* mice phenocopy *Prdm9* null male mice in which sperm production is compromised by inefficient DSB repair and synapsis defects at a pachytene-like stage (Hayashi et al., 2005).

### DSBs are generated at PRDM9-dependent hotspots but are not fully repaired in *Zcwpw1* knockout mice

In PRDM9-null male mice, meiotic DSBs reposition away from PRDM9 bound motifs to promoters and at other sites of PRDM9-independent H3K4me3 (Brick et al., 2012). As *Zcwpw1^KO/KO^* mice phenocopy *Prdm9* null mice, we predicted that DSBs in *Zcwpw1^KO/KO^* mice would be similarly repositioned to promoters. To map DSBs at single base pair resolution, we performed quantitative END-seq (Canela et al., 2016) (Paiano et al., submitted) on testes from *Zcwpw1^KO/KO^* and *Zcwpw1^WT/WT^* juvenile (Figure 6) and adult mice (Figure S6). We identified 2,602 and 3,764 DSB sites in spermatocytes from juvenile *Zcwpw1^WT/WT^* and *Zcwpw1^KO/KO^* mice respectively, and 2,098 and 3,147 DSB sites in spermatocytes from adult *Zcwpw1^WT/WT^* and *Zcwpw1^KO/KO^* mice respectively. Surprisingly, DSBs in both *Zcwpw1^WT/WT^* and *Zcwpw1^KO/KO^* mice nearly completely overlapped with each other and with previously identified hotpots (Figures 6A – 6C and Figures S6A – S6C). Importantly, unlike in *Prdm9* null mice, DSBs were not repositioned towards promoters (Figures 6B and S6B).

**Figure 6:**
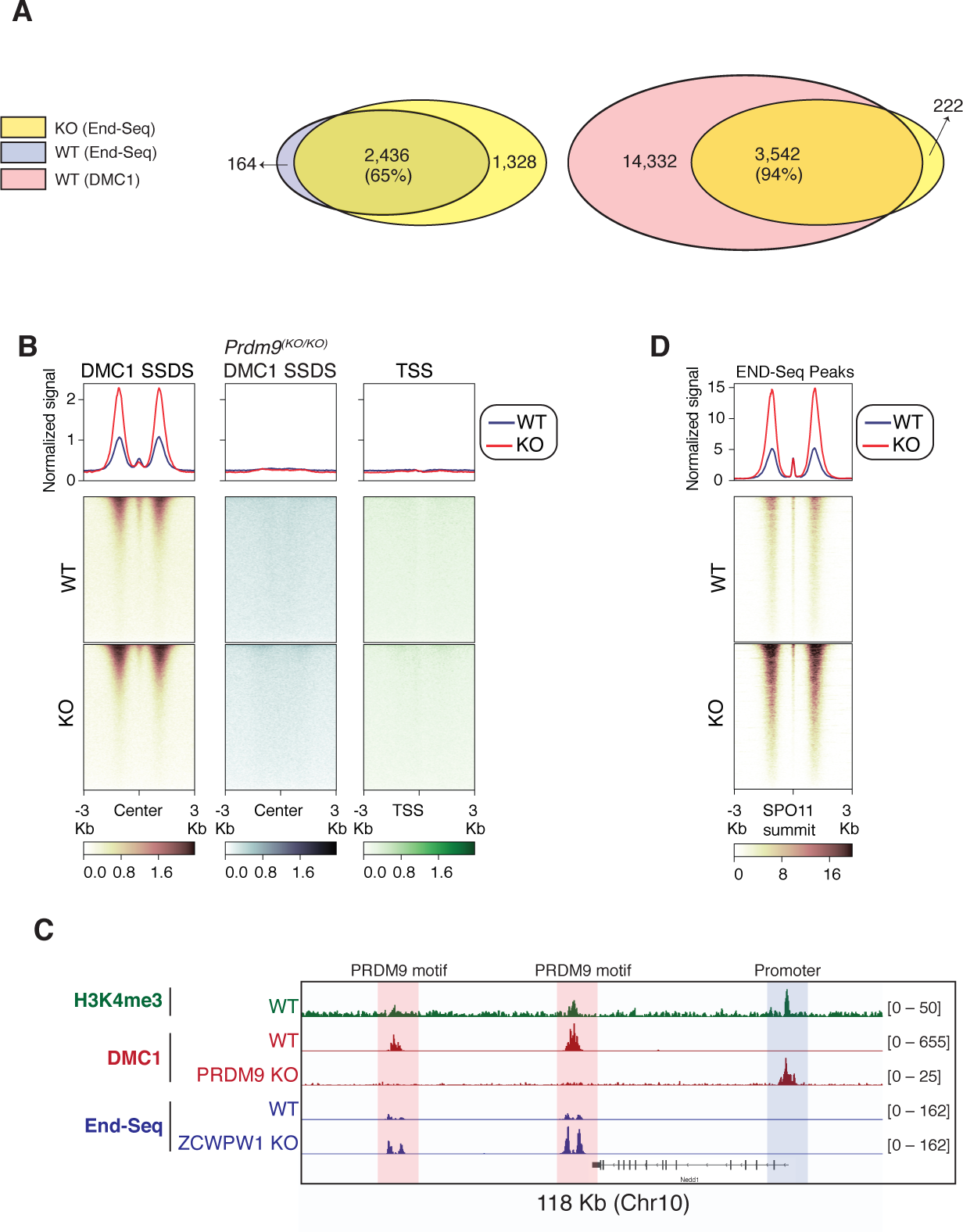
Double Strand Break mapping by END-seq in spermatocytes from juvenile *Zcwpw1^WT/WT^* and *Zcwpw1 ^KO/KO^* mice. (A) Venn diagram showing END-seq peak overlap between juvenile *Zcwpw1^KO/KO^* and *Zcwpw1^WT/WT^* (left) or between *Zcwpw1^KO/KO^* END-seq peaks and DMC1 SSDS hotspots (right). (B) END-seq heatmaps comparing *Zcwpw1^WT/WT^* and *Zcwpw1^KO/KO^* DSBs at different indicated regions: DMC1 SSDS hotspots (WT and PRDM9 KO/ GSE35498) and transcription starting sites (TSS). All signals are normalized with spike-in controls. (C) Read coverage plots for H3K4me3 (CUT&RUN/green), DMC1 SSDS hotspots form WT and PRDM9 KO (GSE35498/red) and END-seq (blue). (D) END-seq aggregate coverage and heatmap plots around SPO11 oligo summits (GSE84689) overlapping with END-seq peaks in WT juvenile mice. Signal is normalized with spike-in controls.

Using spike-in controls to normalize the END-seq signal (Paiano et al., submitted), we estimated an average of 600 breaks per cell in *Zcwpw1^KO/KO^* spermatocytes compared to 402 breaks per cell in *Zcwpw1^WT/WT^* spermatocytes. In addition, after spike-in normalization, we observe a distinct END-seq signal profile across all hotspots, with an increased flanking peak signal with relatively unchanged central peak signal (Figures 6D and S6D). This central peak region largely reflects a recombination intermediate with covalently bound SPO11 resulting from single strand invasion into the unbroken homologous chromosome, whereas the flanking peak regions reflect the extent of end-resection (Paiano et al., submitted). Minimal change in the central peak in comparison to end-resection suggests that, in *Zcwpw1^KO/KO^* spermatocytes, DSB repair is defective downstream of homolog invasion and formation of recombination intermediates. In sum these data indicate that unlike PRDM9, ZCWPW1 is not required for the positioning of DSBs, rather it is required exclusively for efficient DSB repair and later steps of HR.

## Discussion

In this study, we used a computational approach to identify PRDM9 co-expressed genes and identified a novel factor, ZCWPW1, that facilitates the repair of PRDM9-induced DSBs during meiosis. Like PRDM9, ZCWPW1 is expressed at very low/undetectable levels in most tissues but becomes expressed at high levels prior to and during meiotic prophase 1. *Zcwpw1* is necessary for fertility in male mice, as *Zcwpw1^KO/KO^* mice are azoospermic, similar to *Prdm9^KO/KO^*. In contrast, *Zcwpw1* KO females are initially fertile, but suffer from ovarian insufficiency as they age, likely due to delayed meiosis in fetal ovaries (Li et al., 2019a). This is also in contrast to *Prdm9* KO females, which are completely infertile (Hayashi et al., 2005). These data highlight the distinct checkpoint sensitivities in males and females for meiotic progression defects. Furthermore, the distinct phenotypes of *Prdm9* and *Zcwpw1* loss-of-function in females suggest that PRDM9 has ZCWPW1 independent function in females that will require additional exploration.

PRDM9 is a unique SET domain containing histone methyltransferase in several aspects. First it contains its own specific DNA binding domain, a rapidly evolving C2H2 zinc finger array that binds to target sequences with high specificity. Second it contains a dual specificity histone methyltransferase activity for histone H3 K4 and histone H3 K36 (Eram et al., 2014; Powers et al., 2016; Wu et al., 2013). Likewise, ZCWPW1 is a unique protein that possesses dual histone methylation reader domains; a zf-CW domain that has previously been shown to bind to the H3K4me3 mark (He et al., 2010), and a PWWP domain, which has been shown on multiple proteins including DNMT3a and DNMT3b to bind to the H3K36me3 mark (Rondelet et al., 2016). ZCWPW1 can bind to histone H3 peptides with double H3K4me3 and H3K36me3 marks *in vitro* with high affinity at a 1:1 ratio. Furthermore, PRDM9 predominantly methylates H3K4me3 and H3K36me3 on the +1, +2, –1 and –2 nucleosomes flanking its binding sites, which provides a short platform for ZCWPW1 interaction with chromatin at sites in vivo. Thus, we prefer a model in which ZCWPW1 interacts with both marks on the same H3 peptide, but it cannot be ruled out that ZCWPW1 binds to the two marks on separate H3 molecules on the same or adjacent nucleosomes. Resolving these possibilities will require additional structural studies.

Against our expectations, ZCWPW1 is dispensable for initiating DSBs at PRDM9-bound hotspots, as DSBs in *Zcwpw1^KO/KO^* male mice are still located at PRDM9 bound sites. Our data instead suggests that ZCWPW1 is critical for the efficient repair of these PRDM9-dependent breaks. This conclusion is supported by the partial asynaptic pachytene-like meiotic arrest observed in *Zcwpw1^KO/KO^*, which coincides with accumulation of γ-H2AX and DMC1, both characterizing failed DSB repair. The findings that PRDM9 histone methyltransferase activity is essential for DSB formation at PRDM9 binding sites (Diagouraga et al., 2018; Powers et al., 2016) would therefore seem to suggest that a yet to be described PRDM9 histone methyl reader may link PRDM9 to the DSB machinery upstream of ZCWPW1.

In *Zcwpw1^KO/KO^*, the END-seq profile at hotspots demonstrates the presence of a central peak similar to WT mice, indicative of a novel MR intermediate dubbed SPO11-RI (recombination intermediate). Such central peaks are lost in *Dmc1^KO/KO^* mice which fail to undergo strand invasion (Paiano et al., submitted). This indicates that ZCWPW1 facilitates meiotic DSB repair downstream of strand invasion. This coupled with recent reports suggesting that symmetrical PRDM9 binding is critical for MR (Hinch et al., 2019), is most consistent with ZCWPW1 functioning as a PRDM9 specific synapsis factor that is necessary to bind both the cut and uncut ortholog to link PRDM9-bound DNA loops to the SC, to facilitate later steps in MR. But how exactly does ZCWPW1 facilitates such synapsis and repair? A hint may lie within the conserved SCP1 domain found at the N-terminus of ZCWPW1 which shares distant homology to a region of the central axis protein SYCP1 that makes a coiled-coil and facilitates dimerization (Seo et al., 2016). Thus, PRDM9 bound loops could be directly tethered to the SC via ZCWPW1.

In this study we have developed and applied two novel, independent and sensitive methods for mapping meiotic hotspots; END-seq, and ZCWPW1 CUT&RUN. These methods have distinct advantages over previous methods, that include SPO11 oligo sequencing (Lange et al., 2016) and SSDS coupled with DMC1 ChIP-seq (Khil et al., 2012). END-seq is a method of directly sequencing DSBs, and we have applied it to a single mouse testis to map thousands of hotspots. This method could therefore be applied to virtually any organism for mapping hotspots as it does not require the use of antibodies. ZCWPW1 CUT&RUN also maps mouse hotspots, by mapping the occupancy of the factor ZCWPW1 that is directly downstream of PRDM9-induced histone methylation marks using a polyclonal antibody. Importantly, ZCWPW1 occupancy correlates strongly with the strength of hotspots in mice. We applied ZCWPW1 CUT&RUN to 300,00 testicular cells (a tiny fraction from the material obtained from a single adult testis) to sensitively map thousands of mouse hotspots. Thus END-seq provides a greater breadth of possible uses including non-model organisms, whereas ZCWPW1 CUT&RUN provides greater sensitivity with lower cell numbers in mice. With optimizations to both techniques, it is likely they could be pushed to significantly lower cell numbers which would also allow hotspot mapping in females or in species with low meiotic cell numbers.

In yeast, plants and at least some vertebrates (those that lost PRDM9 like canids and birds); meiotic hotspots are located in the nucleosome free regions at gene promoters. However, within species that evolved PRDM9, hotspots are specified by the DNA binding activity of the PRDM9 zinc finger array. Therefore, the emergence of PRDM9 was a landmark that re-shaped patterns of MR during evolution. The PRDM9-derived pattern of hotspot selection provided more flexibility in hotspot evolution compared to the ancestral, fixed pattern of hotspots found at promoters and within genes by decoupling hotspot selection away from functional genetic elements. In this work, we identified ZCWPW1 as an essential factor in the PRDM9 hotspot selection system. This system evolved by co-emergence of (i) a histone writer (PRDM9) which catalyzes formation of H3K4me3 and H3K36me3 dual marks and (ii) a histone methylation reader (ZCWPW1) which recognizes these dual marks to facilitate DSB repair at the PRDM9 bound sites. The presence of these two factors is crucial for hotspot selection and successful meiotic recombination in mice.

## Acknowledgments

We would like to thank Kevin Brick and Florencia Pratto for help with mice phenotyping, CUT&RUN data analysis and for their critical comments and discussion on the manuscript. We thank Gang Cheng for providing the F1 hybrid mouse. We thank Rajan Kumar Choudhary and Charmi Mehta for help with cloning and initial purification. We thank Pedro Rocha and Sarah Frail for providing pA-MN and for their help in CUT&RUN optimization. We also thank Steve Coon and Tianwei Li from the NICHD Molecular Genomics Core for NGS support. This work was supported by grants from the National Institutes of Health 1ZIAHD008933 (TSM), GM114306 (XC) and CPRIT RR160029 (XC).

## Author Contributions

MM and TSM conceived the project. MM identified Zcwpw1 from single cell RNA-seq, generated ZCWPW1 recombinant proteins, screened antibodies, performed CUT&RUN experiments and did phenotyping for *Zcwpw1* KO mice. MM and SR generated and maintained Zcwpw*1* KO mice. MB performed Western blot to detect ZCWPW1 protein expression in different tissues. SP, XZ, and XC performed ZCWPW1 peptide binding assays and affinity measurements using ITC. JP under the supervision of AN developed and performed END-seq for mapping DSBs in meiosis. MM and WW performed data analysis. MM and TSM wrote the manuscript. TSM initiated this collaborative work and along with AN and XC designed the scope of the study.

## Methods

### Experimental Model and Subject Details

#### Mouse models

*Zcwpw1* knockout mouse line (*C57BL / 6N-Zcwpw1^em1(IMPC)Tcp^*) was made as part of the KOMP2-Phase2 project at The Centre for Phenogenomics and was purchased from the Canadian Mouse Mutant Repository. F1 B6/CAST hybrid mouse was generated from mating male CAST (CAST/EiJ) and female B6 (C57BL/6J) (Jackson Laboratory). All experiments were done on juvenile or adult mice (≥2 weeks of age). Mice were sacrificed for testes dissection. Mice breeding, maintenance and experiments were done according to NIH guidance for the care and use of laboratory animals.

### Method Details

#### Protein expression and purification

The mouse zCWPW1 fragment containing the zing finger, CW domain, and PWWP domain (residues 1-440; pXC2085) was cloned into a pet28b vector with an N-terminal 6xHis-SUMO tag and transformed into *E. coli* BL21-Codon Plus (DE3)-RIL (Stratagene). An overnight culture was grown in MDAG media, from which 3 L of LB Broth were inoculated and incubated in shaker at 37°C until A600 of 0.6 was reached. The temperature was then lowered to 16 °C and 30 min later 1 mM ZnCl2 was added with subsequent induction by 0.4 mM isopropyl β-D-1-thiogalactopyranoside (IPTG) 30 min later. Cells were harvested after an overnight growth and immediately suspended in 20 mM Tris (pH 8), 500 mM NaCl, 5 % glycerol (v/v), 0.5 mM Tris(2-carboxyethyl)phosphine hydrochloride (TCEP), and 0.1 mM phenylmethanesulfonyl fluoride (PMSF). Cells were lysed by sonification and clarified by centrifugation 47, 850 x g for 1 hour and passed through 3 µm syringe driven filter unit.

The clarified lysate containing mZCWPW1 was then loaded onto a 5 mL His-Trap HP column (GE Healthcare) using NGC^TM^ chromatography system (BioRad), washed and eluted over a linear gradient from 40 mM to 500 mM Imidazole. Fractions containing zCWPW1 were pooled and subjected to an overnight digest by ULP1 protease (purified in-house) at 4°C to remove the 6xHis-SUMO N-terminal tag. The cleaved zCWPW1 was then diluted to 150 mM NaCl using the aforementioned buffer containing no NaCl. This diluted sample was then loaded onto a 5 mL Hi-Trap Q HP column (GE Healthcare) and eluted over a linear gradient of 150 mM to 1000 mM NaCl with the protein eluting at 225 mM NaCl. Fractions with the highest purity were pooled and concentrated via a 10, 000 MWCO centrifugal filter unit (Millipore) to 4.6 mg/mL and flash frozen in liquid N_2_.

#### Antibody production

Rabbit polyclonal antibody for full-length mouse ZCWPW1 was made by GenScript. Rabbits were immunized with the full-length ZCWPW1 recombinant protein, and serum collected after third immunization. Serum was then affinity purified against the full-length ZCWPW1 protein.

#### Peptide binding experiments

Isothermal Titration Calorimetry (ITC) experiments were conducted at 25 °C with a MicroCal PEAQ-ITC automated system (Malvern). H3(1-18)K4me3, H3(21-44)K36me3, and H3(1-44)K4me3K36me3 peptides were ordered from Biomatik. Binding experiments were performed with protein in the sample cell and peptides were injected into the cell with a syringe. H3(1-18) K4me3 experiments were performed with 358 µM peptide in the syringe, and 50 µM protein in the cell. H3(21-44)K36me3 experiments were performed with 600 µM peptide in the syringe and 50 µM protein in the cell. H3(1-44)K4me3K36me3 experiments were performed with 343 µM peptide in the syringe and 25 µM protein in the cell. The ITC experiments were all conducted with a reference power of 10 µcal/s, with 2.5 µL injections stirred at 750 rpm. The injections were performed over 4 sec and with 200 sec interval to allow for equilibrium to be reached in the buffer consisted of 20 mM Tris (pH 8), 75 mM NaCl, 5 % glycerol (v/v), and 0.5 mM TCEP. Binding constants were obtained by fitting data to the “one set of sites” model in the ITC analysis module. Biotin-Streptavidin Pulldown Assays were conducted using C-terminal biotinylated peptides and streptavidin beads. H3(1-21) and H3(21-44) peptides were ordered from Anaspec, and H3(1-43) peptides were obtained from EpiCypher. Binding reactions consisting of protein (13 µg), biotinylated peptides (0.5 µg), and streptavidin beads were conducted using the binding buffer of 20 mM Tris (pH 8), 75 mM NaCl, 5 % glycerol (v/v), 0.5 mM TCEP and 0.1 % Triton X-100 overnight at 4°C on a tabletop shaker. Samples were then washed 5 times with the binding buffer, resuspended in 10 µL SDS loading dye, and heated at 90°C for 5 min before being run on a stain free gel and imaged by GelDoc imager (BIO-RAD).

#### ZCWPW1 CUT&RUN sequencing

To map ZCWPW1 chromatin binding in spermatocytes, Cleavage Under Targets and Release Using Nuclease (CUT&RUN) was performed in accordance to the previously published protocol (Skene et al., 2018). To prepare single cell suspension, testes were dissected and chopped after tunica removal, then incubated for 30 minutes at 37° C in 3 ml dissociation buffer (DMEM + 2 U/ml Dispase (Worthington) + 250 U/ml Collagenase Type 1 (Worthington) + 50 µg/ml DNAse I (Sigma)). Dissociation enzymes were inactivated by adding DMEM containing 10% FBS. 300,000 cells were used for each CUT&RUN reaction. Initially, cells were washed twice with CUT&RUN wash buffer (20 mM HEPES (pH 7.5), 150 mM NaCl, 0.5 mM Spermidine, 1X cOmplete™ EDTA-free Protease Inhibitor Cocktail (Sigma)). Cells were then resuspended in 1 ml wash buffer, and 10 µl of Concanavalin A–coated beads (Bangs Laboratories) suspended in binding buffer (20 mM HEPES-KOH (pH 7.9), 10 mM KCl, 1 mM CaCl2, 1 mM MnCl2) were added to coat the cells with the beads (Concanavalin A–coated beads were activated by pre-washing twice in binding buffer). Cells-beads mixture were incubated with rotation at room temperature (RT) for 10 minutes in Eppendorf tubes. Thereafter, tubes were placed on magnetic stand until the solution turns clear, and the liquid was discarded. Cells were then resuspended in 50 μl of the antibody buffer (Target-antigen antibody + wash buffer + 0.025% Digitonin + 2 mM EDTA) and incubated at 4° C for 1 hour with rotation. Antibodies’ concentrations used were 10 µg/ml, 2 µg/ml and 10 µg/ml for ZCWPW1, H3K4me3 and GFP respectively. After incubation, cells were washed twice via magnet stand using digitonin buffer (wash buffer + 0.025% digitonin) and resuspended in 50 µl digitonin buffer subsequently. 2.5 µl of Protein A–micrococcal nuclease (pA-MN) fusion protein was then added to the cells (final concentration of ∼700 ng/ml) and mixed with rotation at 4° C for 1 hour. Unbound pA-MN was removed by washing twice using digitonin buffer. Then cells were resuspended in 150 µl digitonin buffer, and incubated on ice for 5 minutes to bring samples’ temperature to 0° C. Thereafter, 3 µl CaCl2 (100 mM stock solution) was added to activate pA-MN micrococcal nuclease activity, and samples were incubated in ice (0° C) for 30 minutes. Subsequently, nuclease reaction was stopped by adding 100 µl of 2X stop buffer (340 mM NaCl, 20 mM EDTA, 4 mM EGTA, 0.05% Digitonin, 100 µg/ml RNase A, 50 µg/ml Glycogen). To release CUT&RUN DNA fragments, samples were incubated for 10 min at 37 °C. Finally, samples were centrifuged at 16,000g for 5 min at 4 °C and were then placed on magnetic stands to collect supernatant (∼ 250 µl) containing released DNA fragments. To extract DNA, 2.5 µl of 10% SDS and 1.875 µl Proteinase K (20 mg/ml) were added to the released DNA, with incubation and shaking at 65° C for 35 minutes. To precipitate DNA, 25 µl 3M Sodium Acetate, 2 µl glycoblue (15 mg/ml) (Thermo Fisher) and 687 µl cold 100% ethanol were added, and samples incubated at -20° C overnight, and then centrifuged in 4° C at maximum speed for 20 minutes. Supernatant was then removed, and DNA pellet rinsed with 70% ethanol, followed by another centrifugation. Ethanol was removed and the pellet resuspended in 10-50 µl TE buffer. DNA concentration was measured by Qubit (Thermo Fisher). For each sample, library prep was done with 3 ng of DNA using SMARTer ThruPLEX DNA-Seq Kit (Takara) according to manufacturer’s protocol. Fourteen cycles of PCR were used in library amplification step, with short elongation time of 10 seconds to selectively amplify for short fragment expected from CUT&RUN. Paired-end sequencing was done using HiSeq 2500.

#### CUT&RUN data analysis

Paired-end reads were mapped to mm10 genome assembly by BWA-MEM version 0.7.17 (Li and Durbin, 2009) using default parameters. Peaks were called by MACS2 version 2.1.2 (Zhang et al., 2008) using default parameters (with exception of the parameter -p 0.001). Read coverage bigwig files were generated by bamCoverage tool of deepTools suite (Ramírez et al., 2014) with the options: --MNase --centerReads. Heatmap plotting was done by computeMatrix and plotHeatmap tools. Integrative Genomics Viewer (IGV) (Robinson et al., 2011) was used for read coverage visualization.

#### ZCWPW1 and PRDM9 motif calling

Motifs calling for ZCWPW1 chromatin bound sites was done by HOMER (Heinz et al., 2010) version 4.10.4. First, ZCWPW1 CUT&RUN reads were mapped to mm10 genome assembly by bowtie2 (Langmead and Salzberg, 2012) version 2.3.5 using the options: --local --very-sensitive-local --no-unal --no-mixed --no-discordant --phred33 -I 10 -X 700. HOMER tag directories were made form mapped bam files by makeTagDirectory command with the option: -tbp 1, and those directories were used to generate homer format peaks by findPeaks command, with the option: - style histone. Motif analysis for ZCWPW1 known motifs and de novo ones was performed by findMotifsGenome.pl command with the options: -size 200 -mask -S 5 -len 14,16,18.

To call motifs of PRDM9, previously published peaks from ChIP-seq of PRDM9^dom2^ and PRDM9^Cast^ alleles (GEO: GSE93955 (Grey et al., 2017)) were used as input for HOMER findMotifsGenome.pl command with the options: -size 200 -mask -S 5 -len 14,16,18

#### ZCWPW1 expression in mouse tissues

To check for differential ZCWPW1 expression in mouse tissues, *Zcwpw1^WT/WT^* or Zcwpw1^KO/KO^ mice were sacrificed and dissected. All isolated tissues were washed in PBS and subsequently dissociated to single cells as described for CUT&RUN. Cells were then lysed at 4°C on gentle nutation for 1h in lysis buffer (50mM Tris HCl pH7.5, 150mM NaCl, 1.5mM MgCl2, 0.2% NP-40, 10% glycerol), supplemented with 1X cOmplete™ EDTA-free Protease Inhibitor Cocktail (Sigma) and 250U/µL benzonase (Sigma). After determining protein concentration, 30µg of protein lysate were used for Western blotting. SDS PAGE was performed using NuPAGE™ 4-12% Bis-Tris Protein Gels in MES buffer (ThermoFisher) and proteins were transferred to nitrocellulose membranes. Membranes were subjected to Ponceau staining to evaluate protein content for each sample and transfer efficiency, blocked for 1h RT in 2% milk in TBST and blotted using 1:1000 custom anti-Zcwpw1 rabbit polyclonal antibody and 1:1000 anti-GAPDH antibody (NB300-327, Novus Biologicals) in 2% milk in TBST, overnight at 4°C on gentle nutation. Proteins were detected by enhanced chemiluminescence using a c600 Azure imager (Azure Biosystems).

#### Haematoxylin and eosin (H&E) staining

Testes were dissected from 8 weeks old adult mice and fixed in 10% formalin. Tissues were embedded in paraffin wax and sectioned at 5 μm thickness, then hematoxylin and eosin (H&E) staining was performed (American Histo Labs).

#### Chromosome spreads from spermatocytes

Testes were dissected and decapsulated from 8 weeks old adult mice in 60 mm dish. 200-800 µl of 100 mM sucrose solution were added, followed by gentle chopping and pipetting to dissociate the tissues, and then passed through 70 µm cell strainer to get single cell suspension. Glass slides were pre-cleaned with isopropanol, and then dipped in fixation buffer (1% paraformaldehyde, 0.15% Triton X100, 0.3 mM NaBorate – pH = 8.5). After slide removal from fixation buffer, while small volume of buffer (∼5 µl) was kept on the slide, ∼20 µl of cells in sucrose were added to the slide while tilting it slowly to spread over the cells on the surface. Slides were then incubated in humid tray for 1 hour at room temperature, followed by air drying. Finally, slides were dipped into 0.04% Photoflo (Kodak), dried and saved at –80° C.

#### Immunofluorescence staining

Chromosome spread slides were thawed from –80° C, washed in PBT (PBS + 0.1% Tween-20) and blocked in blocking solution (PBT + 0.15% BSA) at room temperature for 1 hour. After 3 washes in PBT, primary antibody (diluted in blocking solution, 1/50 for anti-DMC1 and 1/200 for anti-SYCP1, anti-SYCP3 and anti-γ-H2AX) incubated overnight at room temperature in humid chamber. Subsequently, slides washed three times and secondary anti-mouse Alexa Fluor 488 and anti-rabbit Alexa Fluor 555 antibodies (1/200 dilution) were incubated at room temperature in humid chamber for 1 hour. Slides were then washed in PBS, counter stained in DAPI and mounted. A Leica DM6000 B microscope was used for imaging.

### END-seq

#### Mouse testicular cell isolation

Adapted from (Baker et al., 2014). Testes were dissected from 12-14 dpp juvenile or > 8 weeks adult male mice and placed into a 6 cm tissue culture dish containing DMEM. Tunica albuginea were removed under a microscope, and tubules were gently dissociated with forceps and placed into 50 mL tube containing 20 mL DMEM. After tubules settled to the bottom of the tube, DMEM was aspirated and replaced with 20 mL DMEM containing 0.5 mg/mL Liberase TM (Roche, 5401127001) and incubated at 32°C for 15 min at 500 rpm. Tubules were washed once with fresh DMEM, replaced with 20 mL DMEM containing 0.5 mg/mL Liberase TM and 100 U DNase I (ThermoFisher, EN0521), and incubated at 32°C for 15 min at 500 rpm. Tubules were disrupted by gentle pipetting and passed through a 70 μm Nylon cell strainer (Falcon) repeatedly until tissue debris was fully removed. Cells were pelleted at 1500 rpm at 4°C for 5 min and washed with 10 mL DMEM. Cells were filtered through a 40 μm Nylon cell strainer (Falcon) repeatedly until debris was fully removed and pelleted at 1500 rpm at 4°C for 5 min. Cells were resuspended in 1 mL PBS and counted.

#### Embedding cells into agarose plugs

Single-cell suspensions of bulk testicular cells were immediately embedded after isolation into 0.75% agarose plugs. After isolation, cells in 1 mL PBS were diluted to 5-7 million bulk cells/mL PBS and separated into 1 mL of cells per 1.5 mL tube for plug making. Spike-in cells were added at 5% of bulk cell number per tube/plug. Multiple plugs were made per sample if necessary, depending on number of mice and total cell number isolated, processed in the same tube, and DNA later combined after plug melting. A detailed description for embedding cells into agarose plugs and general END-seq procedure can be found in (Canela et al., 2017, 2019). Briefly, agarose embedded cells were immediately lysed and digested with Proteinase K after agarose solidification. Plugs were then washed with TE, treated with RNase, and stored at 4°C for no longer than one week before the next series of enzymatic reactions.

#### Enzymatic reactions

Plugs were treated with sequential combination of Exonuclease VII (NEB) for 1 hr at 37°C followed by Exonuclease T (NEB) for 45 min at 24°C to blunt DNA ends before Illumina adapter ligation (Canela et al., 2019). Subsequent steps of A-tailing, adapter ligation, plug melting, chromatin shearing, and second round of adapter ligation for sequencing were performed exactly as previously described (Canela et al., 2017, 2019).

### END-seq Data Analysis

#### Mapping

Reads were aligned to the mouse (GRCm38p2/mm10) genome using Bowtie version 1.1.2 (Langmead et al., 2009) and 3 mismatches were allowed and the best strata for reads were kept with multiple alignments (-n 3 -k 1 -l 50). Functions ‘‘view’’ and ‘‘sort’’ of samtools (Li, 2011; Li et al., 2009) (version 1.6) were used to convert and sort the mapping output to sorted bam file.

#### Peak calling

Peaks were called using MACS 1.4.3 (Zhang et al., 2008). END-seq peaks were called using the parameters: --shiftsize=1000, --nolambda, --nomodel and --keep-dup = all. Peaks with >2.5 fold-enrichment are kept and those within blacklisted regions (https://sites.google.com/site/anshulkundaje/projects/blacklists) were filtered.

#### Total Break Number Estimation

For estimation of total breaks and comparison between genotypes, we added a spike-in control into END-seq samples which consists of a G1-arrested Ableson-transformed pre-B cell line (*Lig4^-/-^*) carrying a single zinc-finger-induced DSB at the TCRβ enhancer. This site is expected to break in all spike-in cells, which were mixed in at a 5% frequency with bulk testicular cells. END-seq signal was calculated, as RPKM, within ±3kb window around all hotspot centers. Total intensity was divided by the signal around the spiked-in breaks and then divided by 20 since the spiked-in was added at a 1:20 ratio (5%).

### Single-cell PRDM9 co-expression analysis

To identify *Prdm9* co-expressed genes during meiosis at single cell level, a published dataset for single-cell RNA-sequencing (GEO: GSE107644 (Chen et al., 2018)) was analyzed using R version 3.6.1 with the R packages scran version 1.12.1 and scater version 1.12.2. Unique Molecular Identifiers (UMI) counts were normalized, then pair correlation analysis was done to calculate Spearman’s correlation coefficient (Rho) for *Prdm9* co-expression.

### ZCWPW1 evolution analysis

ZCWPW1 orthologs were retrieved and combined from OrthoDB (Kriventseva et al., 2019) and Ensembl (Zerbino et al., 2018) databases. PrositeScan tool (de Castro et al., 2006) was used to analyze domain structure of each ortholog (zf-CW and PWWP domains). To plot consensus sequence of ZCWPW1 domains, amino acid sequences of ZCWPW1 orthologues were aligned using MUSCLE (Edgar, 2004), then amino acid sequences corresponding for each domain were extracted using MEME (Bailey et al., 2009). Consensus motif logos were plotted by WebLogo tool (Crooks et al., 2004).

## Data and Code Availability

All data have been deposited to GEO with the accession number GSE139289.

## Supplemental Information legends

**Figure S1.**
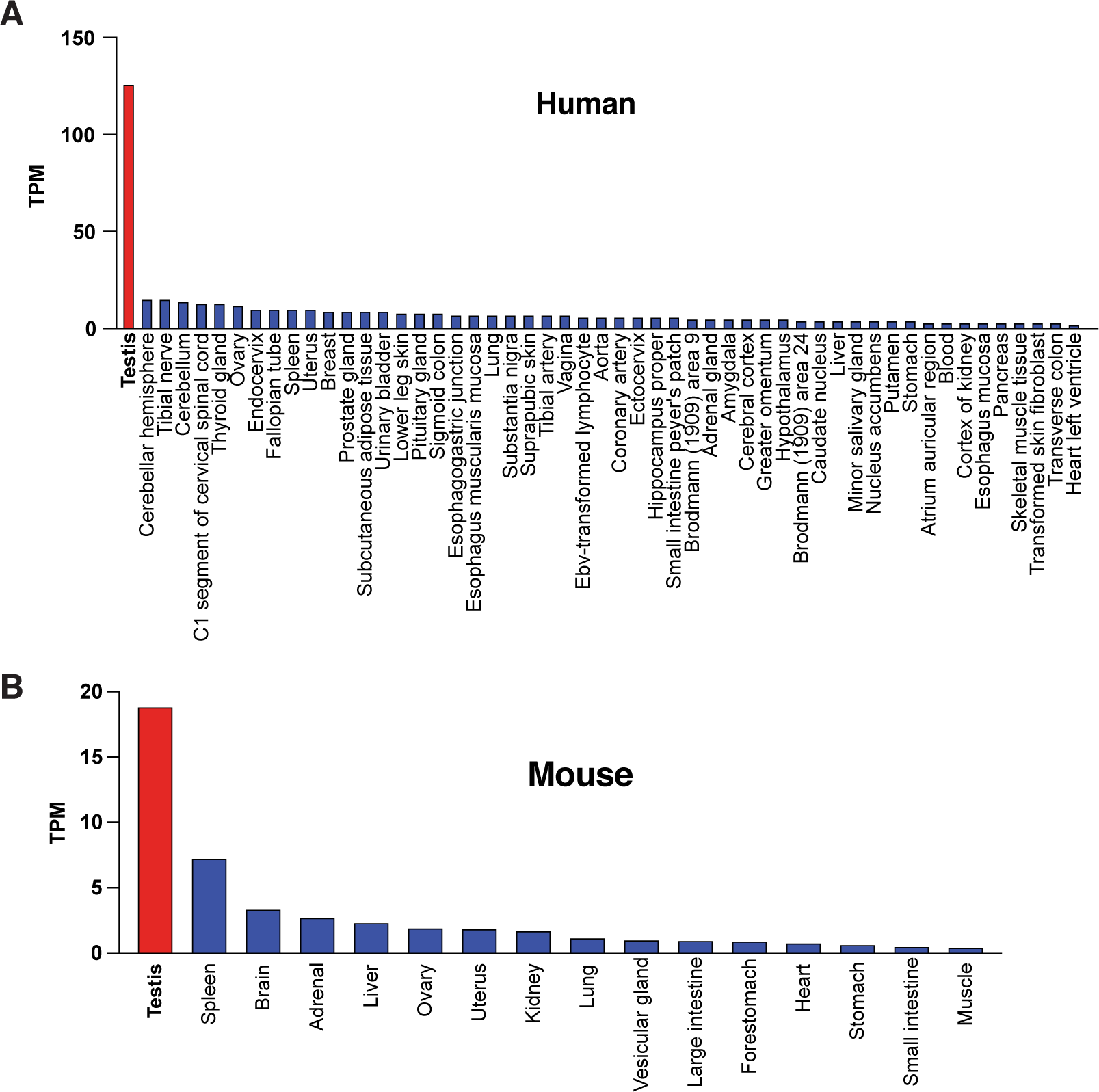
ZCWPW1 tissue expression analysis. Related to Figure 1. (A-B) Tissue distribution of ZCWPW1 transcript expression in human (GTEx) (A), and mouse (Li et al., 2017) (B).

**Figure S2.**
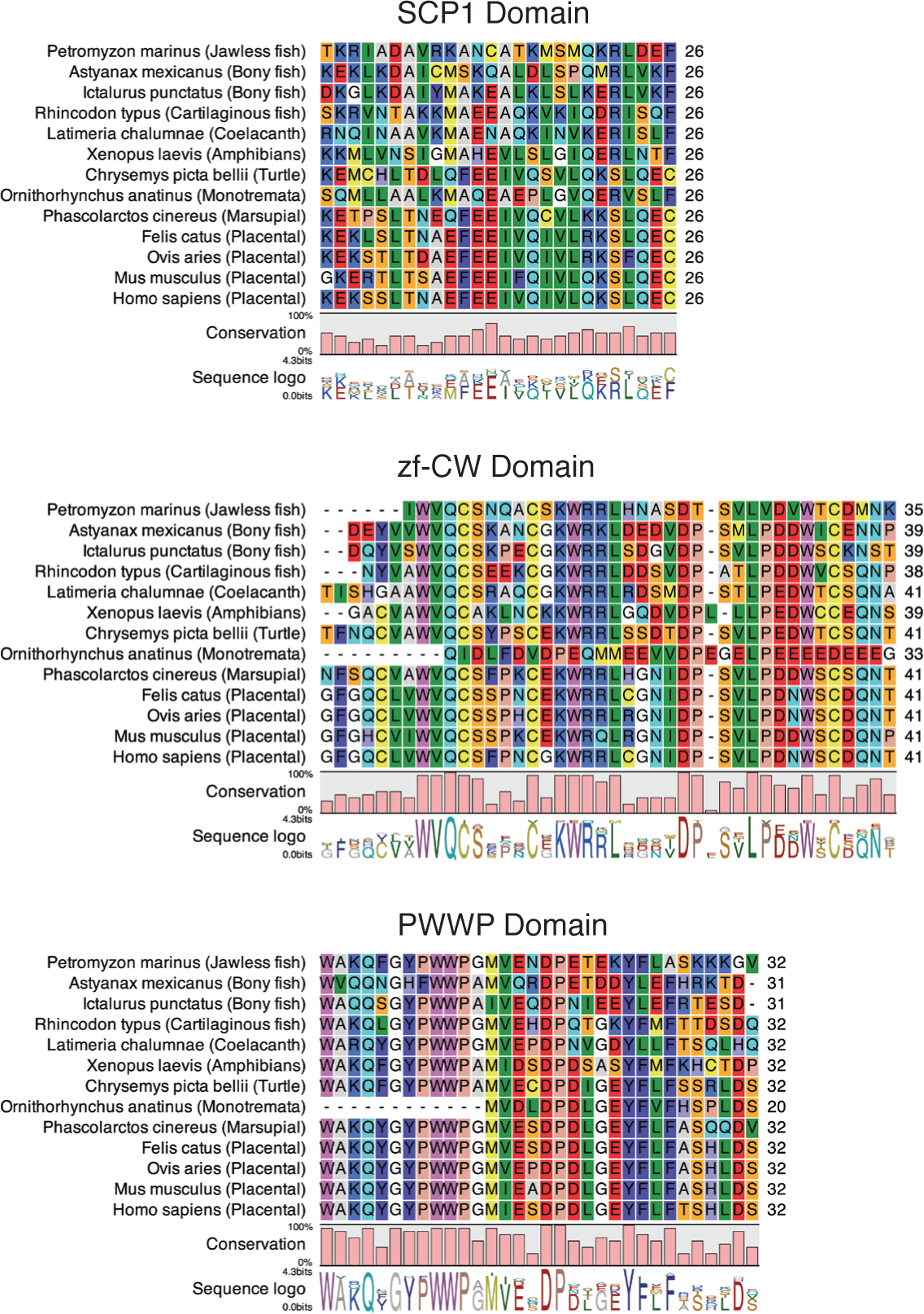
ZCWPW1 multiple sequence alignment of conserved domains. Related to Figure 2. Multiple sequence alignment for regions of conserved domains of ZCWPW1 (SCP1, zf-CW and PWWP) in selected species representative of each clade shown in Figure 2. Alignment for all species is shown in Document S1.

**Figure S3.**
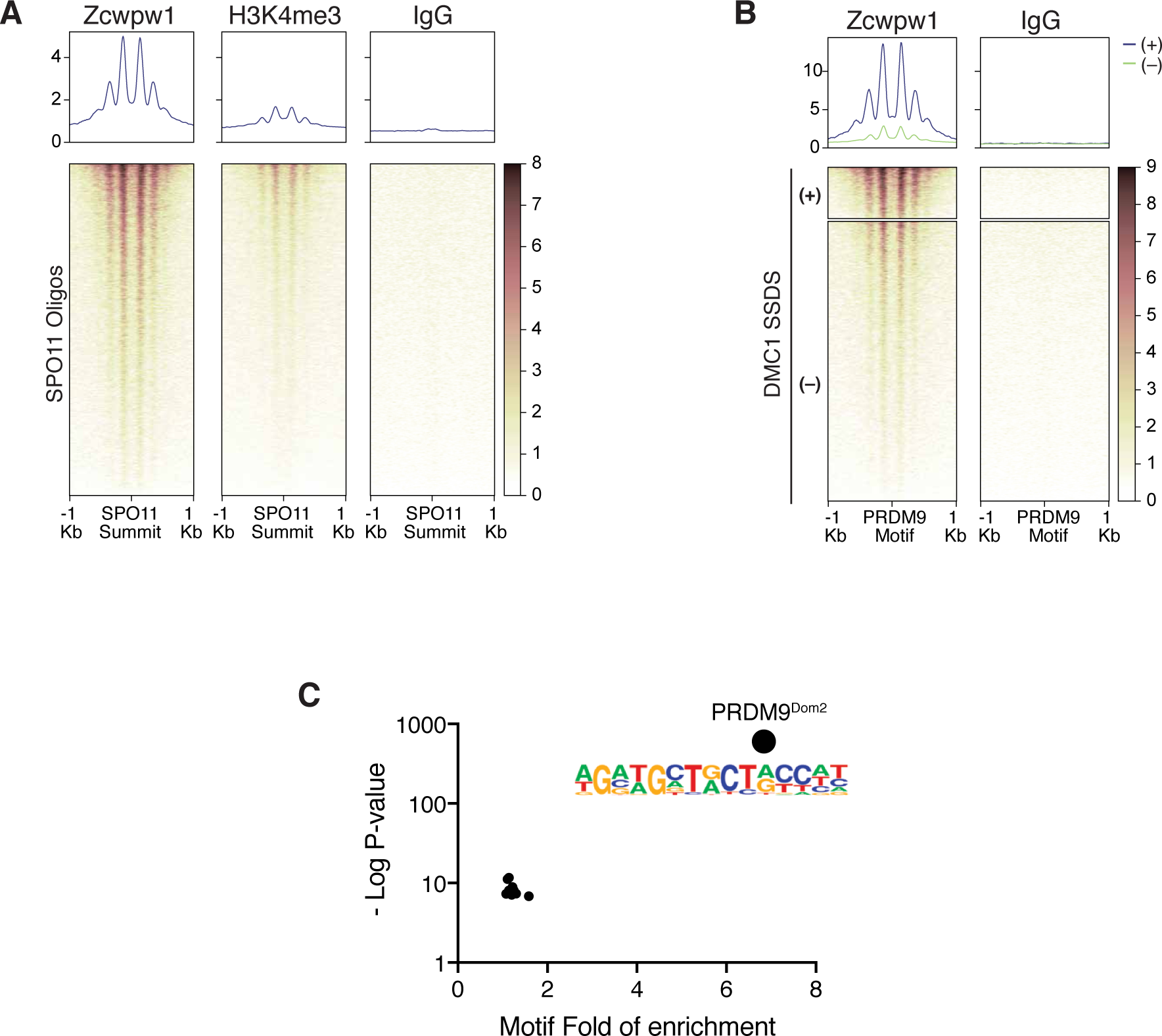
ZCWPW1 chromatin biding *in vivo* in spermatocytes from B6/B6 mice using CUT&RUN. Related to Figure 3. (A) Heatmaps representing ZCWPW1, H3K4me3 and IgG (anti-GFP) occupancy in B6/B6 mice spermatocytes (measured by CUT&RUN) at SPO11 oligos summits (GSE84689) (B) Heatmaps representing ZCWPW1 and IgG (anti-GFP) occupancy in B6/B6 mice spermatocytes at hotspots’ regions determined by DMC1 SSDS (GSE99921). Hotspots were categorized based on their overlap with called ZCWPW1 peaks. (+) refers to hotspots which overlap with ZCWPW1 peak, while (–) refers hotspots which doesn’t overlap with ZCWPW1 peak. Signals are centered around PRDM9 motifs, and any hotspot with multiple motifs was excluded from plotting. (C) Scatter plot for enrichment of all HOMER database known motifs in peaks of ZCWPW1 occupancy in spermatocytes from B6/B6 mice (only motifs with *q* value < 0.05 are shown), logo shown is for HOMER reference PRDM9^Dom2^ motif.

**Figure S4.**
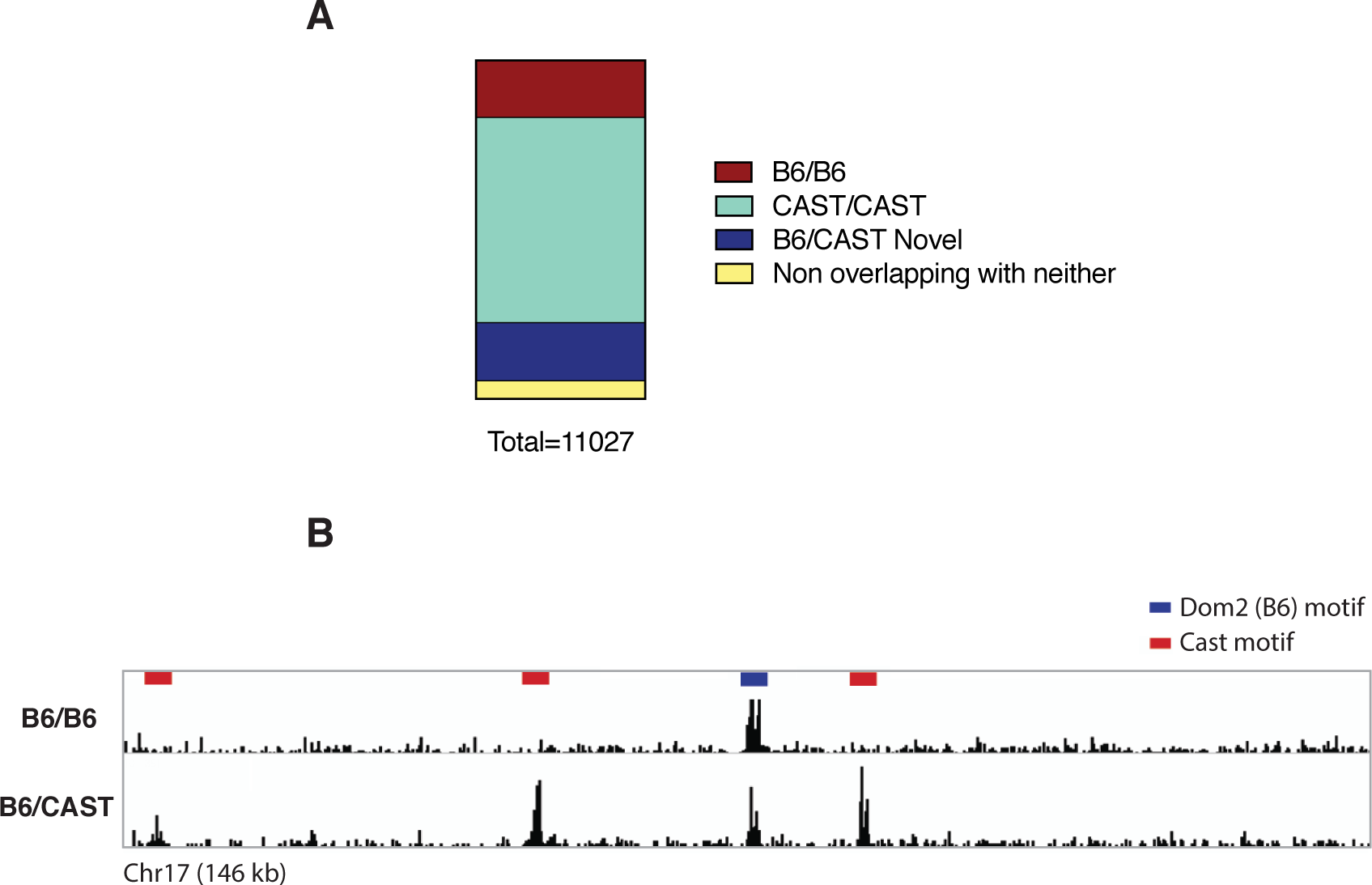
ZCWPW1 chromatin biding *in vivo* in spermatocytes form F1 B6/CAST hybrid mice using CUT&RUN. Related to Figure 4. (A) Chart showing ZCWPW1 peaks overlap with hotspots (DMC1 SSDS / GSE75419) bound by either *Prdm9^Dom2^* or *Prdm9^Cast^* allele. B6/CAST novel hotspots refer to hotspots found in B6/CAST F1 hybrid but not found in either B6/B6 or CAST/CAST spermatocytes. (B) Read coverage plots for ZCWPW1 occupancy in B6/B6 and B6/CAST spermatocytes.

**Figure S5.**
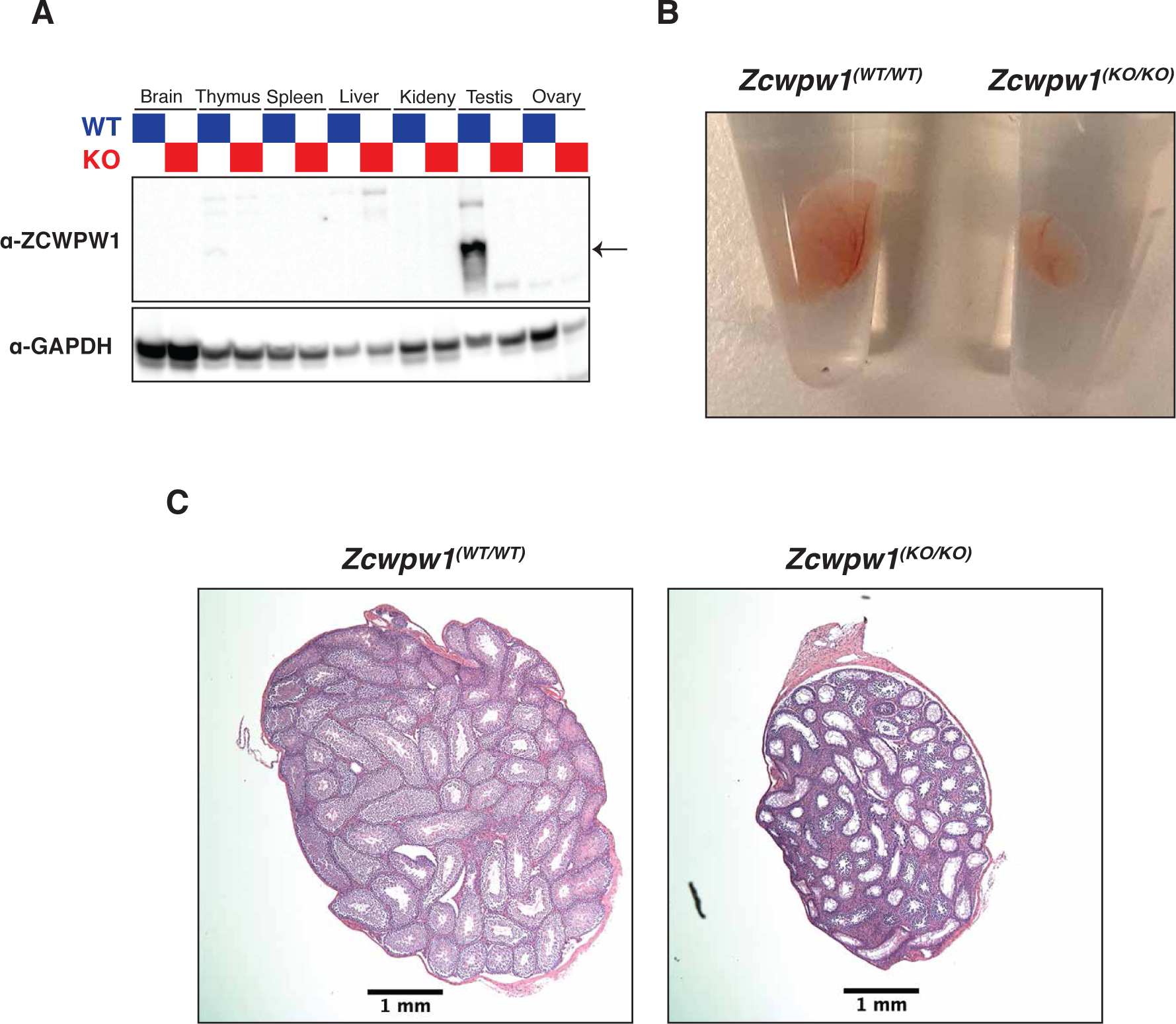
Phenotyping of *Zcwpw1^WT/WT^* and *Zcwpw1^KO/KO^* mice. Related to Figure 5. (A) Western blot with ZCWPW1 specific polyclonal antibody in tissues from WT (blue lanes) or KO mice (red lanes) tissues. The arrow indicates ZCWPW1 band (B) Photo comparing size of testes from *Zcwpw1^WT/WT^* (left) and *Zcwpw1 ^KO/KO^* mice (right) (C) H&E staining of testes sections from *Zcwpw1^WT/WT^* or *Zcwpw1 ^KO/KO^* mice. Scale bar is 1 mm.

**Figure S6.**
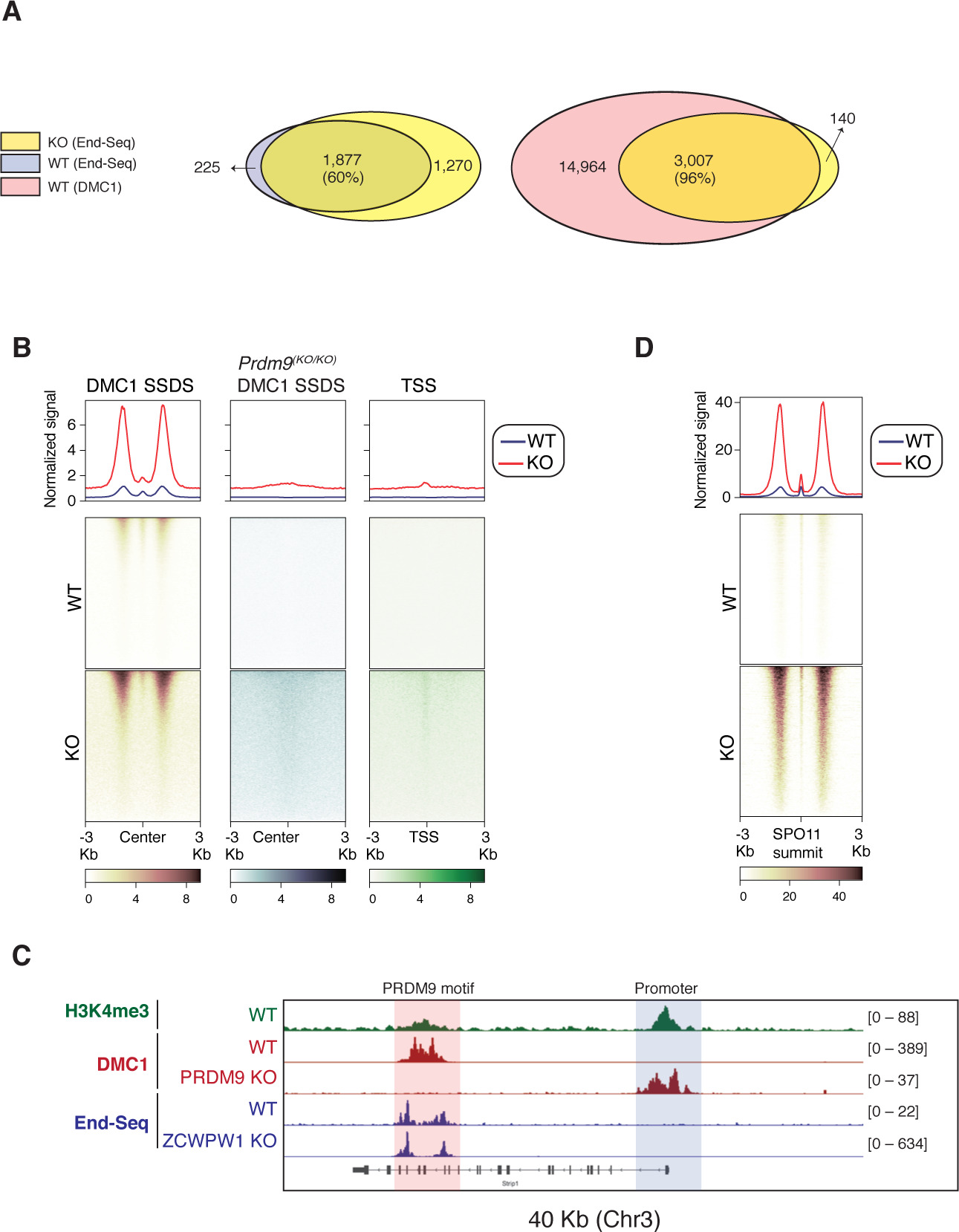
DSBs mapping by END-seq in spermatocytes from adult *Zcwpw1^WT/WT^* and *Zcwpw1 ^KO/KO^* mice. Related to Figure 6. (A) Venn diagram showing END-seq peak overlap between adult *Zcwpw1^KO/KO^* and *Zcwpw1^WT/WT^* (left) and *Zcwpw1^KO/KO^* and DMC1 SSDS hotspots (right) (B) END-seq heatmaps comparing *Zcwpw1^WT/WT^* and *Zcwpw1^KO/KO^* DSBs at different indicated regions: DMC1 SSDS hotspots (WT and PRDM9 KO/ GSE35498) and transcription starting sites (TSS). All signals are normalized to spikes. (C) Read coverage plots for H3K4me3 (CUT&RUN/green), DMC1 SSDS hotspots form WT and PRDM9 KO (GSE35498/red) and END-seq (blue) (D) END-seq aggregate coverage and heatmap plots around SPO11 oligos summits (GSE84689) overlapping with END-seq peaks in WT adult mice. Signal is normalized to spikes.

**Table S1: Prdm9 co-expressed genes during meiosis in single cells. Related to Figure 1.**

Results of Spearman’s correlation analysis between *Prdm9* and each gene in the single cell RNA-seq dataset (Chen et al., 2018)

**Table S2: Zcwpw1 and Prdm9 orthologs. Related to Figure 2.**

List of species for all *Zcwpw1* orthologs retrieved from OrthoDB and Ensembl. Domain structure is based on scanning for presence of zf-CW and PWWP domain in each ortholog using ScanProsite online tool (de Castro et al., 2006). Species from insects (n = 2) and one chordate (Branchiostoma floridae) were excluded from downstream analysis as they lack both domains. List of species for all *Prdm9* orthologs and their domain structure reproduced from (Baker et al. 2017)

**Table S3: Known motifs in ZCWPW1 binding sites. Related to** Figure 3.

Result of screening for enrichment of all known motifs in HOMER database (Heinz et al., 2010). Only motifs with q-value < 0.05 are represented in Figure S3B.

**Document S1: Aligned sequences of SCP1, zf-CW and PWWP domains from all ZCWPW1 orthologs. Related to Figure 2.**

Fasta sequences generated from MEME tool (Bailey et al., 2009) for regions matching ZCWPW1 conserved domains extracted from orthologs sequences. Only orthologs with motif score higher than threshold are included. Full-length protein sequences included in MEME analysis are shown in Table S2.

**Document S2: De novo discovered motifs in PRDM9 and ZCWPW1 bound sites. Related to Figure 4.**

For PRDM9: De novo motifs were called by HOMER from peaks of PRDM9 ChiP-seq dataset GSE93955 (PRDM9^Dom2^ and PRDM9^Cast^) For ZCWPW1: De novo motifs were called by HOMER from peaks of ZCWPW1 CUT&RUN in testes from either B6/B6 or F1 B6/CAST mice.

All motifs are in HOMER motif format. Top motifs are shown in Figure 5A.

