## Supplementary material for "Dual Histone Methyl Reader ZCWPW1 Facilitates Repair of Meiotic Double Strand Breaks": Document S1

**Alignment of SCP-1 motif in ZCWPW1 orthologs**

>Tupaia_chinensis_site_1 offset= 84

KEKSSLTNAEFEEIVQIVLQKSLQECLGM

>Nomascus_leucogenys_site_1 offset= 93

KEKSSLTNAEFEEIVQIVLQKSLQECLGM

>Saimiri_boliviensis_boliviensis_site_1 offset= 91

KEKSSLTNAEFEEIVQIVLQKSLQECLGM

>Sus_scrofa_site_1 offset= 92

KEKSSLTNAEFEEIVQIVLQKSLQECLGM

>Homo_sapiens_site_1 offset= 93

KEKSSLTNAEFEEIVQIVLQKSLQECLGM

>Pongo_abelii_site_1 offset= 93

KEKSSLTNAEFEEIVQIVLQKSLQECLGM

>Gorilla_gorilla_gorilla_site_1 offset= 93

KEKSSLTNAEFEEIVQIVLQKSLQECLGM

>Condylura_cristata_site_1 offset= 92

KEKSSLTNAEFEEIVQIVLRKSLQECLEM

>Marmota_marmota_marmota_site_1 offset= 90

KGKSSLTNAEFEEIVQIVLQKSLQECLGM

>Bubalus_bubalis_site_1 offset= 92

KEKSTLTNAEFEEIVQIVLQKSFQECLGM

>Bos_grunniens_site_1 offset= 94

KEKSTLTNAEFEEIVQIVLQKSFQECLGM

>Vulpes_vulpes_site_1 offset= 82

KEKSSLTNAEFEEIVQIVLRKSLQECLET

>Physeter_catodon_site_1 offset= 93

KGKSTLTNAEFEEIVQIVLQKSLQECLGM

>Tursiops_truncatus_site_1 offset= 92

KGKSTLTNAEFEEIVQIVLQKSLQECLGM

>Orcinus_orca_site_1 offset= 93

KGKSTLTNAEFEEIVQIVLQKSLQECLGM

>Pan_troglodytes_site_1 offset= 93

KEKSSLTNAEFEEIVQTVLQKSLQECLGM

>Pan_paniscus_site_1 offset= 93

KEKSSLTNAEFEEIVQTVLQKSLQECLGM

>Ceratotherium_simum_site_1 offset= 92

KEKSSLTNAEFEEIIQIVLQKSLQECLGM

>Theropithecus_gelada_site_1 offset= 93

KEKSSLTPAEFEEIVQIVLQKSLQECLGM

>Piliocolobus_tephrosceles_site_1 offset= 93

KEKSSLTPAEFEEIVQIVLQKSLQECLGM

>Colobus_angolensis_palliatus_site_1 offset= 93

KEKSSLTPAEFEEIVQIVLQKSLQECLGM

>Rhinopithecus_roxellana_site_1 offset= 93

KEKSSLTPAEFEEIVQIVLQKSLQECLGM

>Rhinopithecus_bieti_site_1 offset= 93

KEKSSLTPAEFEEIVQIVLQKSLQECLGM

>Chlorocebus_sabaeus_site_1 offset= 204

KEKSSLTPAEFEEIVQIVLQKSLQECLGM

>Equus_przewalskii_site_1 offset= 92

KEKSSLTNAEFEEIIQIVLRKSLQECLGM

>Equus_caballus_site_1 offset= 92

KEKSSLTNAEFEEIIQIVLRKSLQECLGM

>Equus_asinus_site_1 offset= 92

KEKSSLTNAEFEEIIQIVLRKSLQECLGM

>Papio_anubis_site_1 offset= 96

KEKSSLTPAEFEEIVQIVLQKSLQECLGM

>Macaca_nemestrina_site_1 offset= 93

KEKSSLTPAEFEEIVQIVLQKSLQECLGM

>Macaca_mulatta_site_1 offset= 96

KEKSSLTPAEFEEIVQIVLQKSLQECLGM

>Macaca_fascicularis_site_1 offset= 96

KEKSSLTPAEFEEIVQIVLQKSLQECLGM

>Cercocebus_atys_site_1 offset= 93

KEKSSLTPAEFEEIVQIVLQKSLQECLGM

>Ictidomys_tridecemlineatus_site_1 offset= 91

KGKSSLTNAEFEEIVQIVLQKSLQECLGT

>Castor_canadensis_site_1 offset= 90

KEKSSLTNAEFEEIVQIVLQKSLQECLGI

>Carlito_syrichta_site_1 offset= 91

KEKSSLTSAEFEEIVQIVLQKSLQECLGM

>Urocitellus_parryii_site_1 offset= 91

KGKSSLTNAEFEEIVQIVLQKSLQECLET

>Spermophilus_dauricus_site_1 offset= 91

KGKSSLTNAEFEEIVQIVLQKSLQECLET

>Bos_indicus_site_1 offset= 94

KEKSTLTNAEFEEIVQIVLQKSFQECLET

>Bos_taurus_site_1 offset= 94

KEKSTLTNAEFEEIVQIVLQKSFQECLET

>Bison_bison_site_1 offset= 93

KEKSTLTNAEFEEIVQIVLQKSFQECLET

>Ursus_maritimus_site_1 offset= 83

KEISSLTNAEFEEIVQIVLRKSLQECLEM

>Callithrix_jacchus_site_1 offset= 95

KEKSSLTNAEFEEIVQIVLQKSLQEHLGM

>Balaenoptera_acutorostrata_site_1 offset= 93

KGKSTLTNAEFEEIVQIVLQKSLQECLET

>Camelus_ferus_site_1 offset= 93

KEKSSLTDAEFEEIVQIVLRKSLQECLET

>Pantholops_hodgsonii_site_1 offset= 92

KEKSTLTDAEFEEIVQIVLRKSFQECLGM

>Vicugna_pacos_site_1 offset= 91

KEKSSLTDAEFEEIVQIVLRKSLQECLET

>Camelus_dromedarius_site_1 offset= 93

KEKSSLTDAEFEEIVQIVLRKSLQECLET

>Camelus_bactrianus_site_1 offset= 93

KEKSSLTDAEFEEIVQIVLRKSLQECLET

>Leptonychotes_weddellii_site_1 offset= 83

KERASLTNAEFEEIVQIVLRKSLQECLEM

>Canis_lupus_familiaris_site_1 offset= 82

KEKSSLTNAEFEEIIQIVLRKSLQECLET

>Acinonyx_jubatus_site_1 offset= 83

KEKLSLTNAEFEEIVQIVLRKSLQECLET

>Panthera_tigris_site_1 offset= 83

KEKLSLTNAEFEEIVQIVLRKSLQECLET

>Panthera_pardus_site_1 offset= 83

KEKLSLTNAEFEEIVQIVLRKSLQECLET

>Felis_catus_site_1 offset= 83

KEKLSLTNAEFEEIVQIVLRKSLQECLET

>Mandrillus_leucophaeus_site_1 offset= 93

KEKSSLTRAEFEEIVQIVLQKSLQECLGM

>Aotus_nancymaae_site_1 offset= 95

KEKSSLTNAEFEEIVQIVLQKSLQERLGM

>Neovison_vison_site_1 offset= 81

KERSSLTSAEFEEIVQIVLQKSLQECLEM

>Odobenus_rosmarus_divergens_site_1 offset= 83

KERASLTNAEFEEIVQIVLRKSLQECLET

>Chrysochloris_asiatica_site_1 offset= 93

KEGASLTNAEFEEIVQIVLQKSLQECLGM

>Ovis_aries_site_1 offset= 92

KEKSTLTDAEFEEIVQIVLRKSFQECLET

>Ovis_aries_musimon_site_1 offset= 93

KEKSTLTDAEFEEIVQIVLRKSFQECLET

>Capra_hircus_site_1 offset= 92

KEKSTLTDAEFEEIVQIVLRKSFQECLET

>Orycteropus_afer_site_1 offset= 94

KERTSLTNAEFEEIVQIVLQKSLQECLGI

>Cebus_capucinus_imitator_site_1 offset= 92

KEKSSLTNAEFEEIVQIVLQKSLQERLGT

>Myotis_lucifugus_site_1 offset= 93

KEKSNLTDEEFEEIVQIVLQKSLQECLGM

>Mustela_putorius_site_1 offset= 82

KERSSLTSAEFEEIVQIVLRKSLQECLET

>Prolemur_simus_site_1 offset= 93

KEKSSISNAEFEEIVQIVLRKSLQECLGM

>Propithecus_coquereli_site_1 offset= 93

KEKSSISNAEFEEIVQIVLRKSLQECLGM

>Lipotes_vexillifer_site_1 offset= 93

KGKSALTNAEFEEIVQIVLQKSLQECLET

>Ailuropoda_melanoleuca_site_1 offset= 83

KEISSITNAEFEEIVQIVLQKSLQECLET

>Neomonachus_schauinslandi_site_1 offset= 96

KERASLTNAEFEEIVQIVLRKSLQECLDT

>Myotis_brandtii_site_1 offset= 93

KEKSNLTNEEFEEIVQIVIQKSLQECLGM

>Eptesicus_fuscus_site_1 offset= 100

KEKSNLTNEEFEEIVQIVIQKSLQECLGM

>Meriones_unguiculatus_site_1 offset= 90

KEKTTLTNAEFEEIFQTVLQKSLQECLET

>Dasypus_novemcinctus_site_1 offset= 93

KEKASLTNAEFEEIVQIVLRKSLQECLEN

>Chinchilla_lanigera_site_1 offset= 86

KEKIILTNAEFEEIVQTVLRKSLQECLGM

>Miniopterus_natalensis_site_1 offset= 93

KEKSSLTNEEFEEIVQIVLQKSLQQCLGI

>Manis_javanica_site_1 offset= 92

KEKASLTNDEFEEIVQIVLQKSLQECLET

>Heterocephalus_glaber_site_1 offset= 91

KEKLILTNAEFEEIVQIVLRKSLKECLGM

>Enhydra_lutris_site_1 offset= 54

KERSSLTSAEFEEIVQIVLRKSLEECLEM

>Delphinapterus_leucas_site_1 offset= 93

KGKSTLTNAEFEELVQIVLQKSLQECLET

>Microcebus_murinus_site_1 offset= 93

KEKSSISNEEFEEIVQIVLRKSLQECLGM

>Pteropus_vampyrus_site_1 offset= 92

KEKSSLTNAEFEEIIQIVLQKSLQQSLGM

>Rousettus_aegyptiacus_site_1 offset= 111

KEKSSLTNAEFEEIIQIVLQKSLQQSLGM

>Choloepus_hoffmanni_site_1 offset= 93

KEKANLTNEEFEEIVQIVLRKSLQECLGI

>Trichechus_manatus_site_1 offset= 93

KEGASLTNAEFEEIVQIVLRKSLQECLGV

>Microtus_ochrogaster_site_1 offset= 83

KEKPTLTNAEFEEIFQIVLQKSLQEYLET

>Ochotona_princeps_site_1 offset= 96

KGRSSLTNAEFEEIVQIVLQKSLQEYEGM

>Fukomys_damarensis_site_1 offset= 98

KEKLILTNAEFEEIVQTVLRKSLKECLGM

>Cavia_porcellus_site_1 offset= 91

KEKLILTNAEFEEIVQTVLRKSLKECLGM

>Loxodonta_africana_site_1 offset= 93

KERGSLTNAEFEEIVQIVLRKSLQECLGS

>Pteropus_alecto_site_1 offset= 92

KEKSSLTNAEFEEIIQIVLQKSLQQSLET

>Odocoileus_virginianus_site_1 offset= 92

KKKSTLTSAEFEEIVQTVLRKSFQECLET

>Myotis_davidii_site_1 offset= 93

KEKSNLTNEEFEEIFQIVIQKSLEECLGM

>Rhinolophus_sinicus_site_1 offset= 93

KEKSSLTNAEFEEIVQLVLQKSLQQSLET

>Cricetulus_griseus_site_1 offset= 90

KEKQTLTNAEFEEIFQIVLRKSLQEYLGL

>Peromyscus_maniculatus_bairdii_site_1 offset= 94

QEKPTLTNAEFEEIFQSVLQKSLQECLGM

>Erinaceus_europaeus_site_1 offset= 91

KEKSSITNEEFEEIVQIVLQKSFQNCLDT

>Jaculus_jaculus_site_1 offset= 93

KGKATLSNAEFEEIVQTVLQKTLQECLGM

>Hipposideros_armiger_site_1 offset= 95

KEKSSLTNAEFEEIFQLVLQKSFQQSLGM

>Mesocricetus_auratus_site_1 offset= 89

KEKQTLTDAEFEEIFQTVLRKSLQEYLET

>Nannospalax_galili_site_1 offset= 100

KEKSTLTNEEFEEIFQTVLKKSLQECLES

>Mus_caroli_site_1 offset= 89

GKEKTLTNAEFEEIFQIVLQKSLQECLET

>Galeopterus_variegatus_site_1 offset= 47

AEKSSLTNEEFEEIVQIVLRKSLQECVAC

>Elephantulus_edwardii_site_1 offset= 90

KERTSLSNAEFEEIVQTVLRKSIEECLET

>Mus_musculus_site_1 offset= 89

GKERTLTSAEFEEIFQIVLQKSLQECLET

>Reference_Domain_SMC_N_site_1 offset= 14

GKERTLTSAEFEEIFQIVLQKSLQECLET

>Rattus_norvegicus_site_1 offset= 88

GKEKTLTSAEFEEIFQIVLQKSLQECLET

>Echinops_telfairi_site_1 offset= 92

KERTSLTNEEYEEIFQIVLQKSREECLET

>Chelonia_mydas_site_1 offset= 100

KEMCHLTDLQFEEIVQCVLQKSLQECMEE

>Oryctolagus_cuniculus_site_1 offset= 96

KGRSSLTNAEFEEIVQTVLRKALQECEAA

>Gopherus_agassizii_site_1 offset= 100

KEMCHLTDLQFEEIVQSVLQKSLQECMEE

>Chrysemys_picta_bellii_site_1 offset= 101

KEMCHLTDLQFEEIVQSVLQKSLQECMEE

>Terrapene_mexicana_triunguis_site_1 offset= 101

KEMCHLTDLQFEEIVQSVLQKSLQECMEQ

>Phascolarctos_cinereus_site_1 offset= 101

KETPSLTNEQFEEIVQCVLKKSLQECLDV

>Mus_pahari_site_1 offset= 18

FWEKTLTNAEYEEIFQTVLQKSLRECLET

>Octodon_degus_site_1 offset= 9

MEKLILTDSEFEEIVQTVLRKSLKECFET

>Vombatus_ursinus_site_1 offset= 101

KETTNLTSEQFEEIVQCVLKKSLQDCLDM

>Pelodiscus_sinensis_site_1 offset= 89

KEMCHLTDLHFEEIVQRALQKSLQECMEE

>Sorex_araneus_site_1 offset= 74

KIKSSLTDEEFEDIVKIVLQKYCQKHLGM

>Pogona_vitticeps_site_1 offset= 71

ERENYLTDAQYDEIFRSVLQKPLQECMEI

**Alignment of zf-CW motif in ZCWPW1 orthologs**

>Enhydra_lutris_site_1 offset= 209

GFGQCLVWVQCSFPNCEKWRRLCGNIDPSVLPDNWSCDQNT

>Chrysochloris_asiatica_site_1 offset= 248

GFGQCLVWVQCSFPNCEKWRRLRGNIDPSVLPDNWSCDQNT

>Condylura_cristata_site_1 offset= 249

GFGQCLVWVQCSFPNCEKWRRLRGNIDPSVLPDNWSCDQNT

>Pteropus_vampyrus_site_1 offset= 249

GFGQCLVWVQCSFPNCEKWRRLRGNIDPSVLPDNWSCDQNT

>Rousettus_aegyptiacus_site_1 offset= 270

GFGQCLVWVQCSFPNCEKWRRLRGNIDPSVLPDNWSCDQNT

>Pteropus_alecto_site_1 offset= 240

GFGQCLVWVQCSFPNCEKWRRLRGNIDPSVLPDNWSCDQNT

>Theropithecus_gelada_site_1 offset= 248

GFGQCLVWVQCSFPNCGKWRRLCGNIDPSVLPDNWSCDQNT

>Piliocolobus_tephrosceles_site_1 offset= 249

GFGQCLVWVQCSFPNCGKWRRLCGNIDPSVLPDNWSCDQNT

>Colobus_angolensis_palliatus_site_1 offset= 249

GFGQCLVWVQCSFPNCGKWRRLCGNIDPSVLPDNWSCDQNT

>Rhinopithecus_roxellana_site_1 offset= 248

GFGQCLVWVQCSFPNCGKWRRLCGNIDPSVLPDNWSCDQNT

>Rhinopithecus_bieti_site_1 offset= 248

GFGQCLVWVQCSFPNCGKWRRLCGNIDPSVLPDNWSCDQNT

>Chlorocebus_sabaeus_site_1 offset= 359

GFGQCLVWVQCSFPNCGKWRRLCGNIDPSVLPDNWSCDQNT

>Acinonyx_jubatus_site_1 offset= 231

GFGQCLVWVQCSSPNCEKWRRLCGNIDPSVLPDNWSCDQNT

>Neomonachus_schauinslandi_site_1 offset= 244

GFGQCLVWVQCSSPNCEKWRRLCGNIDPSVLPDNWSCDQNT

>Panthera_tigris_site_1 offset= 231

GFGQCLVWVQCSSPNCEKWRRLCGNIDPSVLPDNWSCDQNT

>Panthera_pardus_site_1 offset= 231

GFGQCLVWVQCSSPNCEKWRRLCGNIDPSVLPDNWSCDQNT

>Felis_catus_site_1 offset= 231

GFGQCLVWVQCSSPNCEKWRRLCGNIDPSVLPDNWSCDQNT

>Homo_sapiens_site_1 offset= 249

GFGQCLVWVQCSFPNCGKWRRLCGNIDPSVLPDNWSCDQNT

>Pan_troglodytes_site_1 offset= 249

GFGQCLVWVQCSFPNCGKWRRLCGNIDPSVLPDNWSCDQNT

>Pan_paniscus_site_1 offset= 249

GFGQCLVWVQCSFPNCGKWRRLCGNIDPSVLPDNWSCDQNT

>Gorilla_gorilla_gorilla_site_1 offset= 249

GFGQCLVWVQCSFPNCGKWRRLCGNIDPSVLPDNWSCDQNT

>Mandrillus_leucophaeus_site_1 offset= 248

GFGQCLVWVQCSFPNCGKWRRLCGNIDPSVLPDNWSCDQNT

>Papio_anubis_site_1 offset= 252

GFGQCLVWVQCSFPNCGKWRRLCGNIDPSVLPDNWSCDQNT

>Macaca_nemestrina_site_1 offset= 248

GFGQCLVWVQCSFPNCGKWRRLCGNIDPSVLPDNWSCDQNT

>Macaca_mulatta_site_1 offset= 252

GFGQCLVWVQCSFPNCGKWRRLCGNIDPSVLPDNWSCDQNT

>Macaca_fascicularis_site_1 offset= 252

GFGQCLVWVQCSFPNCGKWRRLCGNIDPSVLPDNWSCDQNT

>Cercocebus_atys_site_1 offset= 249

GFGQCLVWVQCSFPNCGKWRRLCGNIDPSVLPDNWSCDQNT

>Camelus_ferus_site_1 offset= 241

GFGQCLVWVQCSSPNCEKWRRLRGNIDPSVLPDNWSCDQNT

>Lipotes_vexillifer_site_1 offset= 242

GFGQCLVWVQCSSPNCEKWRRLRGNIDPSVLPDNWSCDQNT

>Bubalus_bubalis_site_1 offset= 256

GFGQCLVWVQCSSPNCEKWRRLRGNIDPSVLPDNWSCDQNT

>Ictidomys_tridecemlineatus_site_1 offset= 248

GFGQCLVWVQCSSPNCEKWRRLRGNIDPSVLPDNWSCDQNT

>Vicugna_pacos_site_1 offset= 240

GFGQCLVWVQCSSPNCEKWRRLRGNIDPSVLPDNWSCDQNT

>Bos_grunniens_site_1 offset= 258

GFGQCLVWVQCSSPNCEKWRRLRGNIDPSVLPDNWSCDQNT

>Bos_taurus_site_1 offset= 244

GFGQCLVWVQCSSPNCEKWRRLRGNIDPSVLPDNWSCDQNT

>Bison_bison_site_1 offset= 243

GFGQCLVWVQCSSPNCEKWRRLRGNIDPSVLPDNWSCDQNT

>Camelus_dromedarius_site_1 offset= 241

GFGQCLVWVQCSSPNCEKWRRLRGNIDPSVLPDNWSCDQNT

>Camelus_bactrianus_site_1 offset= 241

GFGQCLVWVQCSSPNCEKWRRLRGNIDPSVLPDNWSCDQNT

>Balaenoptera_acutorostrata_site_1 offset= 242

GFGQCLVWVQCSSPNCEKWRRLRGNIDPSVLPDNWSCDQNT

>Physeter_catodon_site_1 offset= 251

GFGQCLVWVQCSSPNCEKWRRLRGNIDPSVLPDNWSCDQNT

>Delphinapterus_leucas_site_1 offset= 242

GFGQCLVWVQCSSPNCEKWRRLRGNIDPSVLPDNWSCDQNT

>Tursiops_truncatus_site_1 offset= 250

GFGQCLVWVQCSSPNCEKWRRLRGNIDPSVLPDNWSCDQNT

>Orcinus_orca_site_1 offset= 249

GFGQCLVWVQCSSPNCEKWRRLRGNIDPSVLPDNWSCDQNT

>Dasypus_novemcinctus_site_1 offset= 243

GFGQCLVWVQCSSPNCEKWRRLRGNIDPSVLPDNWSCDQNT

>Miniopterus_natalensis_site_1 offset= 249

GFGQCLVWVQCSSPNCEKWRRLHGNIDPSVLPDNWSCDQNT

>Odocoileus_virginianus_site_1 offset= 242

GFGQCLVWVQCSSPNCEKWRRLHGNIDPSVLPDNWSCDQNT

>Vulpes_vulpes_site_1 offset= 237

GFGQCLVWVQCSSPNCEKWRRLCGNIDPSVLPDDWSCDQNT

>Galeopterus_variegatus_site_1 offset= 187

GFDQCLVWVQCSFPNCEKWRRLRGNIDPSVLPDNWSCDQNT

>Propithecus_coquereli_site_1 offset= 251

GFGQCLIWVQCSSPNCEKWRRLRGNIDPSVLPDNWSCDQNT

>Neovison_vison_site_1 offset= 237

GFGQCLVWVQCSSPNCEKWRRLCGNVDPSVLPDNWSCDQNT

>Mustela_putorius_site_1 offset= 230

GFGQCLVWVQCSSPNCEKWRRLCGNVDPSVLPDNWSCDQNT

>Sus_scrofa_site_1 offset= 248

GFDQCLVWVQCSSPNCEKWRRLRGNIDPSVLPDNWSCDQNT

>Equus_przewalskii_site_1 offset= 249

GFDQCLVWVQCSSPNCEKWRRLRGNIDPSVLPDNWSCDQNT

>Equus_caballus_site_1 offset= 250

GFDQCLVWVQCSSPNCEKWRRLRGNIDPSVLPDNWSCDQNT

>Equus_asinus_site_1 offset= 249

GFDQCLVWVQCSSPNCEKWRRLRGNIDPSVLPDNWSCDQNT

>Nomascus_leucogenys_site_1 offset= 248

GFGQCLVWVQCSFPNCGKWRRLCGNIDPSVLPDNWYCDQNT

>Ovis_aries_site_1 offset= 241

GFGQCLVWVQCSSPHCEKWRRLRGNIDPSVLPDNWSCDQNT

>Ovis_aries_musimon_site_1 offset= 242

GFGQCLVWVQCSSPHCEKWRRLRGNIDPSVLPDNWSCDQNT

>Capra_hircus_site_1 offset= 242

GFGQCLVWVQCSSPHCEKWRRLRGNIDPSVLPDNWSCDQNT

>Urocitellus_parryii_site_1 offset= 233

GFGQCLVWVQCSSPNCEKWRRLRGNIDPSILPDNWSCDQNT

>Spermophilus_dauricus_site_1 offset= 233

GFGQCLVWVQCSSPNCEKWRRLRGNIDPSILPDNWSCDQNT

>Marmota_marmota_marmota_site_1 offset= 243

GFGQCLVWVQCSSPNCEKWRRLRGNIDPSILPDNWSCDQNT

>Oryctolagus_cuniculus_site_1 offset= 255

GFGQYLVWVQCSSPNCEKWRRLHGNIDPSVLPDNWSCDQNT

>Odobenus_rosmarus_divergens_site_1 offset= 231

GFSQCLVWVQCSSPNCEKWRRLRGNIDPSVLPDNWSCDQNT

>Trichechus_manatus_site_1 offset= 249

GFGQCLVWVQCSSPNCEKWRRLHGNIDPSVLPENWSCDQNT

>Ailuropoda_melanoleuca_site_1 offset= 230

GFGQCLVWVQCSSPDCEKWRRLCGNIDPSVLPDNWSCDQNT

>Cebus_capucinus_imitator_site_1 offset= 248

GFDQCLVWVQCSFPNCGKWRRLCGNVDPSVLPDNWSCDQNT

>Saimiri_boliviensis_boliviensis_site_1 offset= 247

GFDQCLVWVQCSFPNCGKWRRLCGNVDPSVLPDNWSCDQNT

>Aotus_nancymaae_site_1 offset= 251

GFDQCLVWVQCSFPNCGKWRRLCGNVDPSVLPDNWSCDQNT

>Callithrix_jacchus_site_1 offset= 251

GFDQCLVWVQCSFPNCGKWRRLCGNVDPSVLPDNWSCDQNT

>Bos_indicus_site_1 offset= 244

GFGQCLVWVQCSSPNCEKWRRLXGNIDPSVLPDNWSCDQNT

>Pongo_abelii_site_1 offset= 249

GFGQCLVWVQCSLPNCGKWRRLCGNIDPSVLPDNWSCDQNT

>Pantholops_hodgsonii_site_1 offset= 248

GFGQCLVWVQCSSPHCEKWRRLRGNIDPSVLPDDWSCDQNT

>Eptesicus_fuscus_site_1 offset= 251

GFGQCLVWVQCSSPNCEKWRRLRGNIDPSVLPDNWSCVQNT

>Canis_lupus_familiaris_site_1 offset= 230

DFGQCLVWVQCSSPNCEKWRRLCGNIDPSVLPDDWSCDQNT

>Myotis_davidii_site_1 offset= 242

GFGQCLVWVQCSSPNCGKWRRLHGNADPSVLPDNWSCDQNT

>Myotis_brandtii_site_1 offset= 242

GFGQCLVWVQCSSPNCGKWRRLHGNADPSVLPDNWSCDQNT

>Myotis_lucifugus_site_1 offset= 242

GFGQCLVWVQCSSPNCGKWRRLHGNADPSVLPDNWSCDQNT

>Otolemur_garnettii_site_1 offset= 130

GFGQCLIWVQCSSPNCEKWRRLHGNIDPSVLPENWSCDQNT

>Microcebus_murinus_site_1 offset= 251

GFGQCLIWVQCSSPNCEKWRRLRGNIDPAVLPDNWSCDQNT

>Ceratotherium_simum_site_1 offset= 249

GFGQCLVWVQCSSPNCEKWRRLRGNIDPSVLPDSWSCDQNT

>Prolemur_simus_site_1 offset= 249

GFGQCLIWVQCSSTNCEKWRRLRGNIDPSVLPDNWSCDQNT

>Carlito_syrichta_site_1 offset= 244

GFGQCLIWVQCSSPNCEKWRQLRGNIDPSILPDNWSCDQNT

>Loxodonta_africana_site_1 offset= 249

GFGQCLVWVQCSSPNCEKWRRLHRNIDPSALPDNWSCDQNT

>Manis_javanica_site_1 offset= 240

GFGQCQVWVQCSSPNCEKWRRLCGNIDPSVLPVNWSCDQNT

>Ursus_maritimus_site_1 offset= 238

GFGQCLVWVQCSSPDCEKWRRLCGNIDPSVLPANWSCDQNT

>Elephantulus_edwardii_site_1 offset= 229

GFGHCLVWVQCSSPNCEKWRQLRGNIDPSILPENWSCDQNT

>Meriones_unguiculatus_site_1 offset= 234

GFGHCLIWVQCSFPKCEKWRQLRGNIDPSVLPDNWSCDQNP

>Cavia_porcellus_site_1 offset= 249

GFGHCIVWVQCSSPNCRKWRQLCGNIDPSVLPDNWSCNQNT

>Erinaceus_europaeus_site_1 offset= 239

GFGQCQVWVQCSSPNCEKWRRLQGNIDPSVLPDNWSCEQNT

>Microtus_ochrogaster_site_1 offset= 231

GFGQCLIWVQCSFPKCEKWRQLRGDIDPSVLPDNWSCDQNP

>Nannospalax_galili_site_1 offset= 334

GFGQCLVWVQCSFPKCEKWRRLLGNTDPSVLPDNWSCDQNP

>Jaculus_jaculus_site_1 offset= 251

GFGQCIVWVQCSYPTCEKWRRLHRNIDPSVLPDNWSCDQNP

>Octodon_degus_site_1 offset= 159

GFGHCVVWVQCSSPNCRKWRRLCRNIDPSVLPDNWFCHQNT

>Hipposideros_armiger_site_1 offset= 252

GFGQCLVWVQCSFPNCEKWRRLHGNIDPAILPNNWSCEQNT

>Castor_canadensis_site_1 offset= 244

GFGQFLVWVQCSFSNCEKWRRLPGNIDPSVLPDNWSCDQNP

>Ochotona_princeps_site_1 offset= 256

GFGQYLVWVQCSSSNCEKWRRLHGNIDPSVLPDNWSCAQNP

>Mus_musculus_site_1 offset= 239

GFGHCVIWVQCSSPKCEKWRQLRGNIDPSVLPDDWSCDQNP

>Mus_caroli_site_1 offset= 239

GFGHCVIWVQCSSPKCEKWRQLRGNIDPSVLPDDWSCDQNP

>Reference_Domain_zf-CW_site_1 offset= 0

GFGHCVIWVQCSSPKCEKWRQLRGNIDPSVLPDDWSCDQNP

>Peromyscus_maniculatus_bairdii_site_1 offset= 260

GFGQCLIWVQCSSPKCEKWRQLRGGIDPSVLPDNWSCDQNP

>Mesocricetus_auratus_site_1 offset= 238

GFGQCLIWVQCSSAKCEKWRQLRGDIDPSVLPDNWSCDQNP

>Echinops_telfairi_site_1 offset= 241

GFGQYLVWVQCSSPNCEKWRRLCRNMDPSILPSNWSCDQNT

>Cricetulus_griseus_site_1 offset= 252

GFGQCLIWVQCSSSKCEKWRQLRGDIDPSVLPDNWSCDQNP

>Rhinolophus_sinicus_site_1 offset= 248

VTGQCLVWVQCSFPNCGKWRRLRGNIDPSVLPENWSCNQNT

>Mus_pahari_site_1 offset= 168

GFGHCVIWVQCSSPKCEKWRQLRGDIDPSVLPDDWSCDQNP

>Tupaia_chinensis_site_1 offset= 241

GFGLYQVWVQCSFPNCEKWRRLPGNIDPSVLPNNWSCDQNT

>Chinchilla_lanigera_site_1 offset= 237

GFSHCIVWVQCSSPNCRKWRRLCRNIDPSVLPDNWFCYQNT

>Fukomys_damarensis_site_1 offset= 258

GFGHCIVWVQCSFPNCKKWRQLCRNTDPSVLPDNWFCHQNT

>Phascolarctos_cinereus_site_1 offset= 249

NFSQCVAWVQCSFPKCEKWRRLHGNIDPSVLPDDWSCSQNT

>Rattus_norvegicus_site_1 offset= 230

SFGHCVIWVQCSSPKCEKWRQLRGDIDPSVLPDDWSCDQNP

>Vombatus_ursinus_site_1 offset= 249

NFSQCIAWAQCSFPKCEKWRRLHGNIDPSVLPDDWSCSQNT

>Sarcophilus_harrisii_site_1 offset= 51

NFSQCIAWVQCSFPSCAKWRRLLGNTDPSVLPDDWSCSQNT

>Chelonia_mydas_site_1 offset= 242

TFNQCVAWVQCSYPSCEKWRRLSSDIDPSVLPEDWTCSQNT

>Orycteropus_afer_site_1 offset= 171

KTNQFVVWVQCSSPNCEKWRRLHGNIDPSTLPDNWSCAQNI

>Pelodiscus_sinensis_site_1 offset= 231

AFNQCVAWVQCSYPSCEKWRRLSSDVDPSVLPEDWTCSQNT

>Terrapene_mexicana_triunguis_site_1 offset= 243

TFNQCVAWVQCSYPSCEKWRRLSSDTDPSVLPEDWTCSQNT

>Chrysemys_picta_bellii_site_1 offset= 243

TFNQCVAWVQCSYPSCEKWRRLSSDTDPSVLPEDWTCSQNT

>Gopherus_agassizii_site_1 offset= 241

AFNQCVAWVQCSYPSCEKWRRLSSDTDPSVLPKDWTCSQNT

>Notamacropus_eugenii_site_1 offset= 241

XXXQCITWVQCSFPNCEKWRRLHRDIDPSVLPDDWSCSENT

>Chelonoidis_abingdonii_site_1 offset= 104

AVDQCVAWVQCSYPSCEKWRRLSSDTDPSVLPEDWTCSQNT

>Heterocephalus_glaber_site_1 offset= 209

EARPCIVWVQCSSPNCKKWRQLCKNMDPSVLPDNWFCHQNT

>Pogona_vitticeps_site_1 offset= 218

GSCWCIAWVQCSSPTCKKWRQLPSDIDPSVLPEDWSCSQNT

>Thamnophis_sirtalis_site_1 offset= 61

ALGCCTAWVQCSYPSCEKWRRLSSDVDPSALPEDWSCSQNP

>Anolis_carolinensis_site_1 offset= 248

CSSWCIAWIQCSDPNCHKWRRLSSKTDPSVLPEDWSCSQNL

>Ictalurus_punctatus_site_1 offset= 175

DEDQYVSWVQCSKPECGKWRRLSDGVDPSVLPDDWSCKNST

>Astyanax_mexicanus_site_1 offset= 207

ESDEYVVWVQCSKANCGKWRKLDEDVDPSMLPDDWICENNP

>Pygocentrus_nattereri_site_1 offset= 185

DSDKYVVWVQCSRADCFKWRKLSEDVDPSVLPDDWICEDNP

>Latimeria_chalumnae_site_1 offset= 273

TISHGAAWVQCSRAQCGKWRRLRDSMDPSTLPEDWTCSQNA

>Xenopus_laevis_site_1 offset= 127

HRGACVAWVQCAKLNCKKWRRLGQDVDPLLLPEDWCCEQNS

**Alignment of PWWP motif in ZCWPW1 orthologs**

>Vulpes_vulpes_site_1 offset= 310

WAKQYGYPWWPGMVESDPDLGEYFLFASHLDS

>Urocitellus_parryii_site_1 offset= 306

WAKQYGYPWWPGMVESDPDLGEYFLFASHLDS

>Spermophilus_dauricus_site_1 offset= 306

WAKQYGYPWWPGMVESDPDLGEYFLFASHLDS

>Prolemur_simus_site_1 offset= 322

WAKQYGYPWWPGMVESDPDLGEYFLFASHLDS

>Procavia_capensis_site_1 offset= 316

WAKQYGYPWWPGMVESDPDLGEYFLFASHLDS

>Neovison_vison_site_1 offset= 310

WAKQYGYPWWPGMVESDPDLGEYFLFASHLDS

>Nannospalax_galili_site_1 offset= 407

WAKQYGYPWWPGMVESDPDLGEYFLFASHLDS

>Camelus_ferus_site_1 offset= 314

WAKQYGYPWWPGMVESDPDLGEYFLFASHLDS

>Propithecus_coquereli_site_1 offset= 324

WAKQYGYPWWPGMVESDPDLGEYFLFASHLDS

>Hipposideros_armiger_site_1 offset= 325

WAKQYGYPWWPGMVESDPDLGEYFLFASHLDS

>Chrysochloris_asiatica_site_1 offset= 321

WAKQYGYPWWPGMVESDPDLGEYFLFASHLDS

>Pteropus_vampyrus_site_1 offset= 322

WAKQYGYPWWPGMVESDPDLGEYFLFASHLDS

>Lipotes_vexillifer_site_1 offset= 315

WAKQYGYPWWPGMVESDPDLGEYFLFASHLDS

>Myotis_brandtii_site_1 offset= 315

WAKQYGYPWWPGMVESDPDLGEYFLFASHLDS

>Rhinolophus_sinicus_site_1 offset= 321

WAKQYGYPWWPGMVESDPDLGEYFLFASHLDS

>Myotis_lucifugus_site_1 offset= 315

WAKQYGYPWWPGMVESDPDLGEYFLFASHLDS

>Jaculus_jaculus_site_1 offset= 324

WAKQYGYPWWPGMVESDPDLGEYFLFASHLDS

>Ictidomys_tridecemlineatus_site_1 offset= 324

WAKQYGYPWWPGMVESDPDLGEYFLFASHLDS

>Enhydra_lutris_site_1 offset= 282

WAKQYGYPWWPGMVESDPDLGEYFLFASHLDS

>Acinonyx_jubatus_site_1 offset= 304

WAKQYGYPWWPGMVESDPDLGEYFLFASHLDS

>Otolemur_garnettii_site_1 offset= 203

WAKQYGYPWWPGMVESDPDLGEYFLFASHLDS

>Microcebus_murinus_site_1 offset= 324

WAKQYGYPWWPGMVESDPDLGEYFLFASHLDS

>Vicugna_pacos_site_1 offset= 313

WAKQYGYPWWPGMVESDPDLGEYFLFASHLDS

>Neomonachus_schauinslandi_site_1 offset= 317

WAKQYGYPWWPGMVESDPDLGEYFLFASHLDS

>Eptesicus_fuscus_site_1 offset= 324

WAKQYGYPWWPGMVESDPDLGEYFLFASHLDS

>Ursus_maritimus_site_1 offset= 311

WAKQYGYPWWPGMVESDPDLGEYFLFASHLDS

>Elephantulus_edwardii_site_1 offset= 302

WAKQYGYPWWPGMVESDPDLGEYFLFASHLDS

>Cavia_porcellus_site_1 offset= 322

WAKQYGYPWWPGMVESDPDLGEYFLFASHLDS

>Marmota_marmota_marmota_site_1 offset= 316

WAKQYGYPWWPGMVESDPDLGEYFLFASHLDS

>Manis_javanica_site_1 offset= 313

WAKQYGYPWWPGMVESDPDLGEYFLFASHLDS

>Camelus_dromedarius_site_1 offset= 314

WAKQYGYPWWPGMVESDPDLGEYFLFASHLDS

>Camelus_bactrianus_site_1 offset= 314

WAKQYGYPWWPGMVESDPDLGEYFLFASHLDS

>Sus_scrofa_site_1 offset= 321

WAKQYGYPWWPGMVESDPDLGEYFLFASHLDS

>Loxodonta_africana_site_1 offset= 322

WAKQYGYPWWPGMVESDPDLGEYFLFASHLDS

>Trichechus_manatus_site_1 offset= 322

WAKQYGYPWWPGMVESDPDLGEYFLFASHLDS

>Balaenoptera_acutorostrata_site_1 offset= 315

WAKQYGYPWWPGMVESDPDLGEYFLFASHLDS

>Physeter_catodon_site_1 offset= 324

WAKQYGYPWWPGMVESDPDLGEYFLFASHLDS

>Delphinapterus_leucas_site_1 offset= 315

WAKQYGYPWWPGMVESDPDLGEYFLFASHLDS

>Tursiops_truncatus_site_1 offset= 323

WAKQYGYPWWPGMVESDPDLGEYFLFASHLDS

>Orcinus_orca_site_1 offset= 322

WAKQYGYPWWPGMVESDPDLGEYFLFASHLDS

>Odobenus_rosmarus_divergens_site_1 offset= 304

WAKQYGYPWWPGMVESDPDLGEYFLFASHLDS

>Panthera_tigris_site_1 offset= 304

WAKQYGYPWWPGMVESDPDLGEYFLFASHLDS

>Panthera_pardus_site_1 offset= 304

WAKQYGYPWWPGMVESDPDLGEYFLFASHLDS

>Felis_catus_site_1 offset= 304

WAKQYGYPWWPGMVESDPDLGEYFLFASHLDS

>Mustela_putorius_site_1 offset= 303

WAKQYGYPWWPGMVESDPDLGEYFLFASHLDS

>Ailuropoda_melanoleuca_site_1 offset= 303

WAKQYGYPWWPGMVESDPDLGEYFLFASHLDS

>Canis_lupus_familiaris_site_1 offset= 303

WAKQYGYPWWPGMVESDPDLGEYFLFASHLDS

>Rousettus_aegyptiacus_site_1 offset= 343

WAKQYGYPWWPGMVESDPDLGEYFLFASHLDS

>Pteropus_alecto_site_1 offset= 313

WAKQYGYPWWPGMVESDPDLGEYFLFASHLDS

>Dasypus_novemcinctus_site_1 offset= 316

WAKQYGYPWWPGMVESDPDLGEYFLFASHLDS

>Galeopterus_variegatus_site_1 offset= 260

WAKQYGYPWWPGMIESDPDLGEYFLFASHLDS

>Miniopterus_natalensis_site_1 offset= 322

WAKQYGYPWWPGMIESDPDLGEYFLFASHLDS

>Peromyscus_maniculatus_bairdii_site_1 offset= 333

WAKQYGYPWWPGMIESDPDLGEYFLFASHLDS

>Heterocephalus_glaber_site_1 offset= 282

WAKQYGYPWWPGMIESDPDLGEYFLFASHLDS

>Meriones_unguiculatus_site_1 offset= 307

WAKQYGYPWWPGMIESDPDLGEYFLFASHLDS

>Ceratotherium_simum_site_1 offset= 322

WAKQYGYPWWPGMIESDPDLGEYFLFASHLDS

>Equus_przewalskii_site_1 offset= 322

WAKQYGYPWWPGMIESDPDLGEYFLFASHLDS

>Equus_caballus_site_1 offset= 323

WAKQYGYPWWPGMIESDPDLGEYFLFASHLDS

>Equus_asinus_site_1 offset= 322

WAKQYGYPWWPGMIESDPDLGEYFLFASHLDS

>Carlito_syrichta_site_1 offset= 317

WAKQYGYPWWPGMVESDPDLGEYFLFTSHLDS

>Theropithecus_gelada_site_1 offset= 321

WAKQYGYPWWPGMIESDPDLGEYFLFTSHLDS

>Cebus_capucinus_imitator_site_1 offset= 321

WAKQYGYPWWPGMIESDPDLGEYFLFTSHLDS

>Nomascus_leucogenys_site_1 offset= 321

WAKQYGYPWWPGMIESDPDLGEYFLFTSHLDS

>Rhinopithecus_roxellana_site_1 offset= 321

WAKQYGYPWWPGMIESDPDLGEYFLFTSHLDS

>Rhinopithecus_bieti_site_1 offset= 321

WAKQYGYPWWPGMIESDPDLGEYFLFTSHLDS

>Chlorocebus_sabaeus_site_1 offset= 432

WAKQYGYPWWPGMIESDPDLGEYFLFTSHLDS

>Saimiri_boliviensis_boliviensis_site_1 offset= 320

WAKQYGYPWWPGMIESDPDLGEYFLFTSHLDS

>Aotus_nancymaae_site_1 offset= 324

WAKQYGYPWWPGMIESDPDLGEYFLFTSHLDS

>Homo_sapiens_site_1 offset= 323

WAKQYGYPWWPGMIESDPDLGEYFLFTSHLDS

>Pongo_abelii_site_1 offset= 322

WAKQYGYPWWPGMIESDPDLGEYFLFTSHLDS

>Pan_troglodytes_site_1 offset= 323

WAKQYGYPWWPGMIESDPDLGEYFLFTSHLDS

>Pan_paniscus_site_1 offset= 323

WAKQYGYPWWPGMIESDPDLGEYFLFTSHLDS

>Gorilla_gorilla_gorilla_site_1 offset= 323

WAKQYGYPWWPGMIESDPDLGEYFLFTSHLDS

>Mandrillus_leucophaeus_site_1 offset= 321

WAKQYGYPWWPGMIESDPDLGEYFLFTSHLDS

>Papio_anubis_site_1 offset= 326

WAKQYGYPWWPGMIESDPDLGEYFLFTSHLDS

>Macaca_nemestrina_site_1 offset= 321

WAKQYGYPWWPGMIESDPDLGEYFLFTSHLDS

>Macaca_mulatta_site_1 offset= 326

WAKQYGYPWWPGMIESDPDLGEYFLFTSHLDS

>Macaca_fascicularis_site_1 offset= 326

WAKQYGYPWWPGMIESDPDLGEYFLFTSHLDS

>Cercocebus_atys_site_1 offset= 322

WAKQYGYPWWPGMIESDPDLGEYFLFTSHLDS

>Callithrix_jacchus_site_1 offset= 324

WAKQYGYPWWPGMIESDPDLGEYFLFTSHLDS

>Bubalus_bubalis_site_1 offset= 329

WAKQYGYPWWPGMVEPDPDLGEYFLFASHLDS

>Pantholops_hodgsonii_site_1 offset= 321

WAKQYGYPWWPGMVEPDPDLGEYFLFASHLDS

>Bos_grunniens_site_1 offset= 331

WAKQYGYPWWPGMVEPDPDLGEYFLFASHLDS

>Ovis_aries_site_1 offset= 314

WAKQYGYPWWPGMVEPDPDLGEYFLFASHLDS

>Ovis_aries_musimon_site_1 offset= 315

WAKQYGYPWWPGMVEPDPDLGEYFLFASHLDS

>Capra_hircus_site_1 offset= 315

WAKQYGYPWWPGMVEPDPDLGEYFLFASHLDS

>Bos_indicus_site_1 offset= 317

WAKQYGYPWWPGMVEPDPDLGEYFLFASHLDS

>Bos_taurus_site_1 offset= 317

WAKQYGYPWWPGMVEPDPDLGEYFLFASHLDS

>Bison_bison_site_1 offset= 316

WAKQYGYPWWPGMVEPDPDLGEYFLFASHLDS

>Odocoileus_virginianus_site_1 offset= 315

WAKQYGYPWWPGMVEPDPDLGEYFLFASHLDS

>Ochotona_princeps_site_1 offset= 329

WAKQYGYPWWPGMVESDPDLGEYFLFASHLDL

>Condylura_cristata_site_1 offset= 322

WAKQYGYPWWPGMVESDPDLGEYFLFSSHLDS

>Mesocricetus_auratus_site_1 offset= 311

WAKQYGYPWWPGMIETDPDLGEYFLFASHLDS

>Mus_pahari_site_1 offset= 241

WAKQYGYPWWPGMIEADPDLGEYFLFASHLDS

>Mus_musculus_site_1 offset= 312

WAKQYGYPWWPGMIEADPDLGEYFLFASHLDS

>Mus_caroli_site_1 offset= 312

WAKQYGYPWWPGMIEADPDLGEYFLFASHLDS

>Reference_Domain_PWWP_site_1 offset= 14

WAKQYGYPWWPGMIEADPDLGEYFLFASHLDS

>Fukomys_damarensis_site_1 offset= 331

WAKQYGYPWWPGMIESDPDLGEYFLFASHLHS

>Erinaceus_europaeus_site_1 offset= 312

WAKQHGYPWWPGMVESDPDLGEYFLFASHLDS

>Tupaia_chinensis_site_1 offset= 314

WAKQYGYPWWPGIIESDPDLGEYFLFASHLDS

>Myotis_davidii_site_1 offset= 315

WAKQYGYPWWPGMVELDPDLGEYFLFASHLDS

>Oryctolagus_cuniculus_site_1 offset= 328

WAKQYGYPWWPGMIESDPDLGEYFLFASQLDS

>Cricetulus_griseus_site_1 offset= 325

WAKQYGYPWWPGMIETDPDLGEYFLFSSHLDS

>Echinops_telfairi_site_1 offset= 314

WAKQYGYPWWPGMVEPDPDLGEYFLFASHLDC

>Microtus_ochrogaster_site_1 offset= 304

WAKQHGYPWWPGMIEPDPDLGEYFLFASHLDS

>Octodon_degus_site_1 offset= 232

WAKQYGYPWWPGMVESDPDLEEYFLFASHFDS

>Rattus_norvegicus_site_1 offset= 303

WAKQYGYPWWPGMIECDPDLGEYVLFASHLDS

>Piliocolobus_tephrosceles_site_1 offset= 323

WAKQYGYPWWPGVIECDPDLGEYFLFTSHLDS

>Colobus_angolensis_palliatus_site_1 offset= 322

WAKQYGYPWWPGVIECDPDLGEYFLFTSHLDS

>Chinchilla_lanigera_site_1 offset= 310

WAKQYGYPWWPGMVEADPDLEEYFLFASHFDS

>Vombatus_ursinus_site_1 offset= 322

WAKQYGYPWWPGMVESDPDLGEYFLFASQQDV

>Phascolarctos_cinereus_site_1 offset= 322

WAKQYGYPWWPGMVESDPDLGEYFLFASQQDV

>Sarcophilus_harrisii_site_1 offset= 124

WAKQYGYPWWPGMVESDPDLGEYFLFASQQDV

>Pelodiscus_sinensis_site_1 offset= 304

WAKQFGYPWWPGMVECDPDIGEYFLFSSRLDS

>Gopherus_agassizii_site_1 offset= 314

WAKQFGYPWWPAMVECDPDIGEYFLFSSRLDS

>Chelonoidis_abingdonii_site_1 offset= 177

WAKQFGYPWWPAMVECDPDIGEYFLFSSRLDS

>Terrapene_mexicana_triunguis_site_1 offset= 316

WAKQFGYPWWPAMVECDPDIGEYFLFSSRLDS

>Chrysemys_picta_bellii_site_1 offset= 316

WAKQFGYPWWPAMVECDPDIGEYFLFSSRLDS

>Chelonia_mydas_site_1 offset= 315

WAKQFGYPWWPAMVECDPDIGEYFLFSSRLDS

>Thamnophis_sirtalis_site_1 offset= 134

WAKQYGYPWWPGIIEADPDIGEYFLFSSQADS

>Pogona_vitticeps_site_1 offset= 291

WAKQYGYPWWPGLIEADPDIEEYFLFSSQMDL

>Anolis_carolinensis_site_1 offset= 321

WAKQYGYAWWPGVIEADPYLGEYLLFSSQTDS

>Latimeria_chalumnae_site_1 offset= 346

WARQYGYPWWPGMVEPDPNVGDYLLFTSQLHQ

>Notamacropus_eugenii_site_1 offset= 314

WAKQHGYPWWPGMIESDPDLEQYFLFGSPQEP

>Sorex_araneus_site_1 offset= 228

EGSGFGFQRWPGMVESDPDLGEYFLFASHLDS

>Orycteropus_afer_site_1 offset= 221

EEKKIREIQWPGMVESDPDLGEYFLFASHLDS

>Rhincodon_typus_site_1 offset= 70

WAKQLGYPWWPGMVEHDPQTGKYFMFTTDSDQ

>Dipodomys_ordii_site_1 offset= 95

GEGRIDFSWWPGMVESDPDLGEYFLFASHLDF

>Xenopus_laevis_site_1 offset= 199

WAKQFGYPWWPAMIDSDPDSASYFMFKHCTDP

>Xenopus_tropicalis_site_1 offset= 32

WAKQSGYPWWPAMIDSDPDSAYFFKFKHCTDP

>Pygocentrus_nattereri_site_1 offset= 258

WAWQSGHPWWPAMIERDPDTYDYLEFHRKTDL

>Oncorhynchus_tshawytscha_site_1 offset= 281

WAQQIGYPWWPAIVERDPDTKTFCQFNRNTDL

>Salmo_salar_site_1 offset= 182

WAQQIGYPWWPAIVERDPDTKTFCQFNRNTDL

>Oncorhynchus_mykiss_site_1 offset= 334

WAQQIGYPWWPAIVERDPDTKTFCQFNRNTDL

>Petromyzon_marinus_site_1 offset= 65

WAKQFGYPWWPGMVENDPETEKYFLASKKKGV

>Lepisosteus_oculatus_site_1 offset= 460

WARQFGYPWWPAMVEKDPETGDYMEFQNSRIS

>Ictalurus_punctatus_site_1 offset= 248

WAQQSGYPWWPAIVEQDPNIEEYLEFRTESDL

>Astyanax_mexicanus_site_1 offset= 280

WVQQNGHFWWPAMVQRDPETDDYLEFHRKTDL
